# A joint distribution framework to improve presence-only species distribution models by exploiting opportunistic surveys

**DOI:** 10.1101/2021.06.28.450233

**Authors:** Juan M. Escamilla Molgora, Luigi Sedda, Peter Diggle, Peter M. Atkinson

**Affiliations:** Lancaster Environment Centre, Lancaster University, Lancaster LA14YQ, UK; Centre for Health Informatics, Computing and Statistics (CHICAS), Lancaster Medical School, Faculty of Health and Medicine, Lancaster University, Lancaster LA1 4YQ, UK; Lancaster Medical School, Faculty of Health and Medicine, Lancaster University, Lancaster LA1 4YQ, UK; Faculty of Science and Technology, Lancaster University, Lancaster LA1 4YR, UK

**Keywords:** species distribution models, presence-only data, opportunistic sampling, multivariate conditional autoregressive models, model-based statistical ecology

## Abstract

**Aim:** We propose a Bayesian framework for modelling species distributions using presence-only biodiversity occurrences obtained from historical opportunistic surveys.

**Location:** Global applicability with two case studies in south-east Mexico.

**Methods:** The framework defines a bivariate spatial process separable into ecological and sampling effort processes that jointly generate occurrence observations of biodiversity records. Presence-only data are conceived as incomplete observations where some presences have been filtered out. A choosing principle is used to separate out presences, missing data and absences relative to the species of interest and the sampling observations. The framework provides three modelling alternatives for accounting the spatial autocorrelation structure: independent latent variables (model I); common latent spatial random effect (model II); and correlated latent spatial random effects (model III).

The framework was compared against the Maximum Entropy (MaxEnt) algorithm in two case studies: one for the prediction of pines (Class: Pinopsida), using botanical records as sampling observations and another for the prediction of Flycatchers (Family: Tyranidae), using bird sightings as sampling records.

**ăResults:** In both case studies, at least one of the proposed models achieved higher predictive accuracy than MaxEnt. The model with correlated spatial effects fit best when the sampling effort was informative, while the one with a shared spatial effect was more suitable in cases with high proportion of non sampled sites.

**Main Conclusions:** Our approach provides a flexible framework for presence-only SDMs aided by a sampling effort process informed by the accumulated observations of independent and heterogeneous surveys. For the two case studies, the framework provided a model with a higher predictive accuracy than an optimised version MaxEnt.

## 1. Introduction

Species distribution models (SDMs) are statistical and computational methods for characterising the distribution of organisms across space (Guisan and Zimmermann, 2000; Elith and Leathwick, 2009). The predictive capabilities of these models allow forecasting changes in species distribution under different environmental scenarios, providing meaningful insights in which to assess biodiversity loss (Pereira et al., 2010), adaptation to climate change (Wiens et al., 2009), ecosystem management and conservation (Navarro et al., 2017) or risk of invasive species (Jiménez-Valverde et al., 2011). Modelling species distributions have helped to develop strategies for management, adaptation and mitigation of human-induced impacts to the biosphere (Ferrier et al., 2016; Foden and Young, 2016; Intergovernmental Panel on Climate Change, 2014).

SDMs use occurrence observations as response variable(s) and environmental features (covariates) as explanatory variables. The methodological frameworks for estimating species distributions are diverse. For example, early methods for estimating potential distributions include the method of environmental envelope (Booth, 1985) for characterising suitability areas correlated with climatic variables. Later, generalised linear models (GLMs) and generalised additive models (GAMs) (Guisan and Zimmermann, 2000; Guisan et al., 2002) and (Keating and Cherry, 2004) were used to model distributions based on presence and absence records. Machine learning methods have also been used. Specifically, supervised classification algorithms have been extensively used (e.g Segurado and Araújo (2004); Elith et al. (2006); Peterson et al. (2011)). These methods include boosted regression trees (BRT, Friedman (2001)), multivariate adaptive regression spline (MARS, Friedman (1991)) and artificial neural networks (ANN, Rosenblatt (1958)). The R package `sdm` (Naimi and Araújo, 2016) includes an exhaustive list of machine learning methods for fitting species distribution models.

One of the main concerns in applying machine learning methods for predicting species distributions is the abstraction of complex ecological processes into a black-box classification machine that does not explicitly describe the stochastic nature that generates the observations, limiting their scientific interpretability (Haegeman and Loreau, 2008; Gelfand and Shirota, 2019). In this sense, model-based statistical methods are better fit to describe the underlying mechanisms of species distributions. In particular, joint stochastic modelling and hierarchical Bayesian models have recently been proposed to account for uncertainties in the parameters estimations and for defining more flexible random effects. For example, in cases where spatial autocorrelation is present, the use of Gaussian Processes (Golding and Purse, 2016) or Gaussian Markov Random Fields (GMRF) (Illian et al., 2013) have been shown to increase predictive accuracy. Although these models are statistically sound, their major limitation is their reliance on presence-absence data, which generally are not available. In cases where the goal is the modelling of species distributions across large geographic regions, the collection of presence-absence records requires a careful sampling design with possibly hundreds of experts deployed in the field for data collection. Surveys of this kind are atypical and usually are developed by governments or similar sized institutions that can afford full inventory or census data (e.g. forest Inventory and analysis (Smith, 2002) and Inventario Nacional Forestal (CONAFOR, 2018)).

The widespread use of opportunistic observations has been favoured by citizen science initiatives and the availability of large and open repositories like: The Global Biodiversity Information Facility GBIF (GBIF Secretariat, 2015), eBird for bird sightings (Hudson et al., 2014) and the PREDICTS database (Sullivan et al., 2009)). These records are often derived from museums, herbaria collections or unstructured citizen observations. As such, the data are often limited to presence-only observations and, therefore, do not include information on where or when a given species was *not* found (i.e. absences). In addition, the information related to sampling design is frequently lost, or does not exist, and the data itself are prone to several sources of bias in space, time, and detectability among species and habitats (Dickinson et al., 2010; Beck et al., 2014; Isaac and Pocock, 2015; Franklin et al., 2016). Despite the inevitable problem of their sampling bias, presence-only observations contain valuable information about species distributions and, therefore, several modelling frameworks for presence-only data have been proposed for such purposes.

With the exception of some unrealistic assumptions about the absences on presence-only models (e.g. assuming that absence of evidence is equivalent to evidence of absence), estimating the probability for species occurrence using solely presence-only observations involves a problem of model identification (Ward et al., 2009). That is, the model has multiple solutions and is not possible to make reliable inferences. This problem has lead to recognise the importance of incorporating other sources of information into SDMs based on presence-only data.

One of the earliest methods is the Maximum Entropy (MaxEnt) algorithm (Phillips et al., 2006) for predicting occurrences based on the density of environmental covariates conditional to the known species presences using background data. The background data are samples from the available area and can include presences or absence of observations. The MaxEnt algorithm reduces predictions to an optimal density distribution calculated with a constrained optimization algorithm, ignoring accountability for uncertainties related to the optimised distribution and the specification of other random effects. Despite this, it has shown to perform well in practice (Elith et al., 2006) and is still one of the most widely used methods for predicting species distributions (> 2600 articles in Web of Science at the time of writing).

Phillips et al. (2009) recognised the effect of the sampling bias in presence-only distribution models and proposed the use of occurrence records of other species that are have been collected using the similar methods (called a “target group” in the sense of Phillips et al. (2009)). In their work, they proposed a joint model for accounting the sampling bias and implemented their methodology in three generic types of models: GAMs, MARS, BRTs and Maxent. Their conclusion was that using and informed background data (one that potentially shares same characteristics of the sampling process) significantly improves the models’ accuracy.

The use of joint modelling methods for accounting sampling bias has been addressed by other authors. For example, the expectation maximization algorithm for estimating underlying presence-absence processes (Ward et al., 2009) aims to infer the underlying presence-absence logistic signal of the data used as presence-only observations. This approach does not account for spatial dependencies. The occupancy model proposed by Royle and Kéry (2007) specifies a hierarchical Bayesian model for accounting the joint effect of two components, one for imperfectly observed occupancy and the other for detections conditional on that process. Inconveniently, this particular model is suited for longitudinal data (i.e. time series) and does not account for any spatial effect.

In this regard, the framework developed by Pacifici et al. (2017) accounts spatial dependencies in both components, one for presence-only data and other based on presence-absence. However, both proposals do not allow the explicit modelling of the preferential sampling.

Although these models have advanced the SDMs in many aspects, a more integrated spatial statistical framework for species distributions using presence-only data that can explicitly model the spatial influence of the sampling effort is still needed. We consider that a framework of this kind with the capability for jointly modelling the sampling effort and the ecological processes using a flexible design for defining missing data can contribute to a greater predictive accuracy by exploiting citizen science effort.

We present a statistical framework for modelling species distributions using presence-only data. We assume that the registered occurrences of a taxon of interest (ToI) are incomplete observations of a bivariate process that includes information about the environmental suitability (i.e. where the ToI can live) and complementary occurrence data that serve as a proxy for sampling effort, providing information on how the observations were recorded. The framework specifies three hierarchical bayesian models that jointly specifies the ecological and sampling processes. The approach provides a full description of the data generating process, giving a more direct interpretation of the parameters as well as giving explicit estimates of their uncertainties. The presented model assumes that the species populations are static in time and in equilibrium with the environment (in the sense of Guisan and Zimmermann (2000)). Therefore, this model does not differentiate between sink populations or populations with sustained growth.

The paper is structured as follows. Section 2 describes the general specification of the frameworks. Here, we develop a logistic hierarchical model defined as a bivariate process that accounts for spatial random effects. Our most general model (full description in appendix: Appendix A.3.3) includes a latent bivariate spatial process with correlated components. We also consider two extreme special cases: in model I (appendix: Appendix A.3.1) the two component processes are independent; in model II (appendix: Appendix A.3.2) they are proportional. In section 3 we propose two study cases for predicting presences of Pines (class: *Pinopsida*) and Flycatchers (family: *Tyrannidae*). The prediction analysis is described in sections 4.1 and 4.2, respectively. We compared the framework using the three models with the MaxEnt algorithm as a standard benchmark. Finally, section 5 discusses the methodology, caveats and future research.

## 2. Materials and Methods

As presence-only data lack real absences, there exists no knowledge on whether the absence of data is due to the inaccessibility of a potential sampling location or the real absence of the taxon of interest (ToI). This ambiguity suggests that presence-only data provide incomplete evidence of two underlying processes acting together. A process *P_Y_* that generates the ecological phenomenon of a taxon’s occurrence, and a process *P_X_* associated with the sampling effort or survey. As such, locations with no records of the ecological phenomenon or sampling effort indicates incomplete or missing information. Our proposal is an attempt to model these two processes using a hierarchical Bayesian framework with the aim to predict probability of occurrence for a ToI using presence-only data under different configurations of the spatial autocorrelation of *X* and *Y*.

### 2.1. Model summary

In general, the framework specifies a Bayesian hierarchical model that accounts for the joint effect of two components; an ecological process (*P_Y_*), that drives the occurrence of species of interest in the study region, and a sampling effort process (*P_X_*) that models how the occurrence data were sampled. Each stochastic process include a structural component (fixed effect) and a random effect that includes the specification of spatial autocorrelation. The model is defined in a discrete spatial lattice. Consequently the estimations are also discrete and are defined in each area element of the lattice. The support of the model is the area element.

The presence-only data is assumed to represent realizations of a bivariate stochastic binary process (Bernoulli) separable in two components: one relative to an ecological process *P_Y_* that drives the environmental suitability for the ToI, and another process *P_X_* related to the sampling effort. *P_X_* and *P_Y_* are modelled according to the following equations:

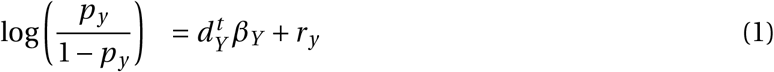

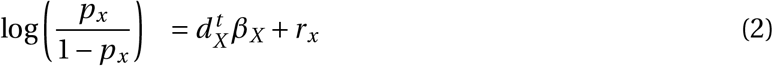

where *d_X_* and *d_Y_* represent vectors of explanatory variables and *r_X_* and *r_Y_* the random effects for *X* and *Y*, respectively. Specifically, *d_Y_* is suited for environmental variables of ecological importance, while *d_X_* should account for variables that help explain the sampling process.

The data used to fit both processes includes information on known occurrences of the ToI, the sampling effort and missing observations. To predict the probability for sites with missing data, we use the *data augmentation* scheme proposed by Tanner and Wong (1987) and implemented by Lee (2013) in the R-Cran package *CARBayes*. The approach generates posterior samples of *X* and *Y* as well as the latent variables related to processes *P_Y_* and *P_X_* in all locations, including the ones with missing observations (i.e. 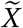 and 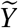).

The full model specification is explained in the supplementary materials Appendix A.

#### 2.1.1. Three models for spatial variation

The proposed framework assumes that the ecological process *P_Y_* and the anthropogenic sampling process *P_X_* are conditionally independent given the random effects *R_Y_* and *R_X_*. Figure 1 show the model structure while a detailed description of the framework specification is in the supplementary materials Appendix A.

**Figure 1:**
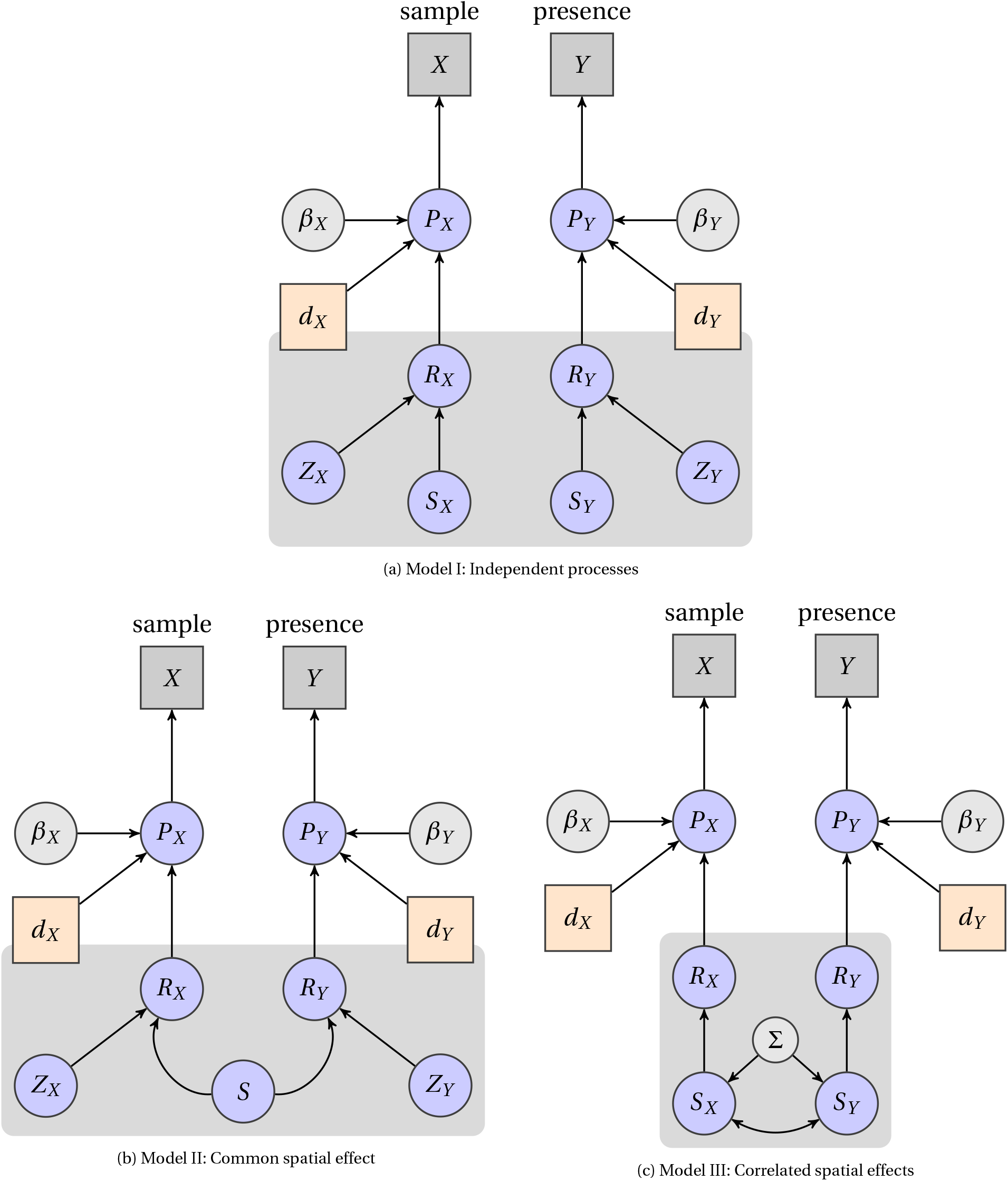
Directed acyclic graphs for the three model specifications. Variables in squares account for observations: *Y*: presence of a taxon of interest (e.g. species) and *X*: presence of sample. Circles in blue correspond to latent variables while circles in grey correspond to parameters. Variables *P_X_* and *P_Y_* correspond to the latent processes of the sampling effort and environmental suitability, variables *R_X_* and *R_Y_* correspond to the random effect for the sampling effort and the environmental suitability processes respectively. Variables *β_x_* and *β_Y_* represent the parameters of the fixed effects (linear components) of the latent processes *P_X_* and *P_Y_* respectively. Squares in salmon colour indicate environmental (*d_Y_*) and anthropic (*d_X_*) explanatory variables. The variables inside the dark grey block define the random effects component; different in the three models. Variables *S*, *S_X_* and *S_Y_* describe the spatial component defined as Gaussian Markov Random Fields, while variables *Z_X_* and *Z_Y_* represent unstructured variability within an area.

The spatial random effect are described by components *S_Y_* (ToI) and *S_X_* (sampling effort). The only source of dependency between *R_Y_* and *R_X_* is the dependency between these spatial components. In addition, each random effect incorporates an independent component for modelling unstructured variation, namely variables *Z_Y_* and *Z_X_*, corresponding to *R_Y_* and *R_X_* respectively. The framework assumes that the observations of presence for the ToI and the existence of the survey (sampling) are independent when conditioned to the spatial effect. As such, the spatial autocorrelation structure is responsible for informing both processes. To test for this effect we designed three possible models in which the spatial processes *S_Y_* and *S_X_* inform *R_Y_* and *R_X_*. Model I where *S_Y_* and *S_X_* are independent, model II with one shared spatial process (*S_X_* = *S_Y_*) and model III where *S_X_* and *S_Y_* are correlated components. Schematics of the directed acyclic graphs (DAG) describing the three models are reported in figure 1, while the full description of the framework is described in supplementary materials Appendix A.

We are aware that estimating real probability of occurrence using presence-only data is not possible given the inherently sampling bias of these type of data (e.g Guillera-Arroita et al. (2014)). Along this text, we refer to *environmental suitability* as the spatial variation across space that determines a species to live, settle or occupy a given area. This definition disregards the scale of the given value for a particular area. In other situations, we use the term *probability of occurrence* to account for the spatial variation of the ecological process (i.e. environmental suitability) in a probabilistic context, that is, where the spatial variation ranges in values from 0 to 1. To exemplify this compare the range in values of the latent variable *S_Y_* (spatial effect) to those of the ecological process *P_Y_*. Values in *P_Y_* are range only within the [0,1] interval.

#### 2.1.2. Selection of explanatory variables

Our framework is based on the Grinnellian definition of ecological niche, that is, a niche defined by non-interactive and non-consumable (scenopoetic) variables with environmental conditions changing smoothly and coarsely in space (Soberón, 2007). The selection of these explanatory variables (covariates) are crucial for the interpretability of the model and, although, the general specifications for *P_X_* and *P_Y_* are mathematically similar (eqs. A.7 and A.8), they describe very different processes. *P_Y_* models the environmental suitability for a ToI to occupy the area under study. Therefore, its associated explanatory variables (*d_Y_*) should be of ecological interest. Examples of these variables are: temperature, precipitation, evapotranspiration, elevation, slope and vegetation cover. On the other hand, *P_X_* models the probability of a ToI to be sampled, given that it has been observed. This process is assumed to be independent from the environmental suitability and it is fully determined by anthropic variables such as: distance to closest road, population density, infrastructures, political borders or land use type. The selection of covariates depends on the nature and specificities of each problem and research question. Therefore, the classification between anthropic and ecological variables is not necessarily mutually exclusive.

### 2.2. A Choosing Principle for obtaining presences, relative absences and missing observations

Estimating the probability of occurrence using solely presence-only observations necessarily requires additional assumptions about non-existent absences (Ward et al., 2009). Thus, any non recorded presence of the taxon of interest (ToI) can potentially be a real absence (i.e. the area is not inhabited by the ToI) or an unobserved presence (i.e. the ToI inhabits the area but there is not record about it). The fundamental concept of this work is to use occurrence records of other taxa that are considered to share a similar sampling pattern as the ToI. These occurrences are used to model a sample effort process that informs about the presence and absence of the taxon of interest.

Models I, II and III specify a joint bivariate process that uses two vectors of observations as inputs; one (*Y*) for fitting the ecological process (*P_Y_*) and other (*X*) for fitting the associated sampling effort process (*P_X_*). These input vectors (hereafter called *response vectors*) are composed of *k* entries, one for each area element of the spatial lattice. Each entry has assigned one of three possible states: *presence* (1), *relative absence* (0) or *missing data* (N.A). As such, for a given site (*k*), a state of *presence* indicates that the taxa of interest (ToI) has been observed. A state of *relative absence* (0) indicates that the surrogate taxon is present (i.e *X_k_* = 1) but the ToI is absent (i.e. *Y_k_* = 0). A state of *missing data* (also called *missing observations*) indicates that the neither the ToI nor the surrogate taxa are present in the site *k* (i.e. *X_k_* = 0 = *Y_k_*).

As we are using exclusively occurrence data we need an algorithm for deriving response vectors *X* and *Y* from presence-only records. We call this algorithm the *choosing principle* and receives two lists as inputs: *target* 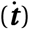 and *background* 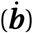. These lists are obtained by checking the existence of an occurrence on each area element of the spatial lattice. That is, if on a given area, there exists at least one record inside, assign a 1, otherwise assign a 0. This procedure is repeated on all the *k* areas of the spatial lattice. Contrary to the response vectors *X* and *Y*, where each entry can be either 1, 0 or N.A., the entries of 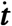 and 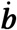 are composed binary (i.e. 0 or 1). Obtaining the missing values (N.A.) is performed by transforming 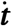 and 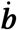 into response vectors *X* and *Y* using the *choosing principle*. As such, the choosing principle defines the missing data for *X* and *Y*, given the presence-absence lists of the target and background observations.

There are many possibilities to define a choosing principle. Here, we used one that, for a given site *i*, assigns: missing data (N.A.) where neither the background nor target observations are present (i.e ***t**_i_* = 0 = ***b**_i_*), 0 where there is no presence of a target observation but has a background observation (i.e. ***t**_i_* = 0 and ***b**_i_* = 1), and 1 to locations where there is presence of the target taxa (i.e ***t**_i_* = 1) Algorithm 1 describes this *choosing principle*.

It is worth noting that, for each response vector, a target 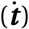 and background 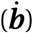 lists are needed. Specifically, for obtaining the response vector of the ToI (*Y*) the target and background list would correspond to the occurrences list of the taxon of interest and the surrogate taxa (or taxon) respectively. In the case of the sample observations (*X*), the target list would correspond to the surrogate taxa while the background list could be any taxonomic group that, upon consideration of the researcher, informs the sampling effort process. A pragmatic selection would be the use of all available records, disregarding their taxonomic classification.

The selected choosing principle is reasonable from an ecological view. If, on average, the existence of *X* informs the occurrence of *Y*, we can argue that: if a site *i* has no background information, the probability of *X* and *Y* is unknown and it is informed only by nearby sites. If on the other hand, the background information exists, but there is no known occurrence (i.e. a *relative absence*) of *Y* at area *i*, the probability of occurrence for *Y* will depend on the presence of *X* as well as its nearby areas. In this sense, the probability of occurrence of a taxon (e.g. species) depends on the presence, its relative absence, its sampling effort and the nearby areas where the taxon is present. The next section shows two practical examples.

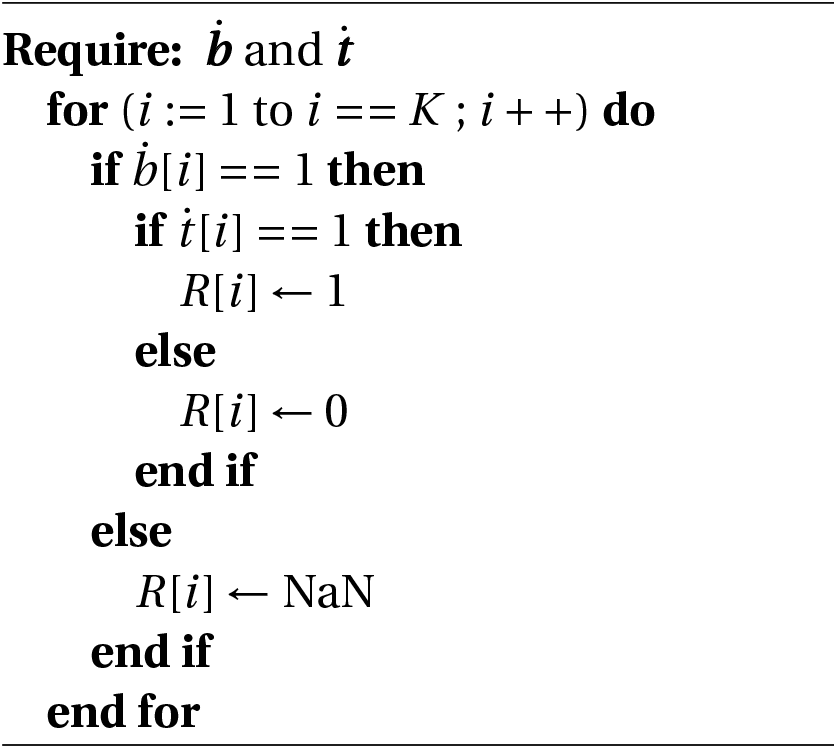
Choosing principle. Obtaining a response vector *R* using background 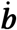 and target observations 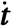 over a spatial lattice composed of *K* area elements. Binary values are: 1 if there is at least one registered occurrence, and 0 otherwise. The symbol *N.A* (*Not a number*) is assigned to missing values.

## 3. Applications

To show the capabilities of the framework we chose two examples for predicting presences. The first involves predicting the presence of pines, that is, occurrences of the class *Pinopsida* as the process *P_Y_* (*Pines*) using the available botanical records and occurrences of the kingdom *Plantae* as the sampling process *P_X_* (*Plants*). The second example predicts the presence of a relatively abundant family of flycatchers (family: *Tyrannidae*) as the process *P_Y_* (*Tyranids*), using the available records of birds (class *Aves*) as the sampling process *P_X_* (*Birds*). In both cases we chose *Elevation* and *Precipitation* as the scenopoetic variables for process *P_Y_* and *Distance to roads* and *Population density* as the anthropological variables for process *P_X_*. Following the model specification in equations A.7 and A.8 (supplementary materials Appendix A) The model for the examples of *Pines* and *flycatchers* is defined as the joint Bernoulli process.

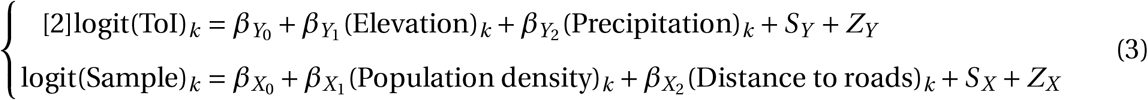

Where the word *ToI* indicates that the equation is used for the taxon of interest (i.e. pines or flycatchers) and *Sample* indicates that the equation is valid for the sampling effort (i.e. plants or birds).

### 3.1. Study region

Both models were fitted to data from the same study region. The region comprises the inland area of a circular polygon centered in central-eastern Mexico at 19N-97E with radius of 2° (ca.~ 200 km). The area covers approximately 112,000 km^2^ and intersects several Mexican states including: Veracruz, Puebla, Tlaxcala, Hidalgo, Mexico City, Morelos and Oaxaca (see figure 2 (i)). It includes heterogeneous landscapes with variability in biodiversity, geomorphological and climatic features. The region also includes distinct biomes such as: coastal dunes, chaparrales, mesophyl forests, evergreen rainforest, grasslands, mangroves, broad leaf forests and coniferous forests (Rzedowski, 2006) and (INEGI, 2015). The circular polygon was intersected on a grid of 4 km spatial resolution to obtain a lattice 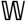 composed of 4061 areal units. This lattice was used to define the spatial structure in models I, II and III.

**Figure 2:**
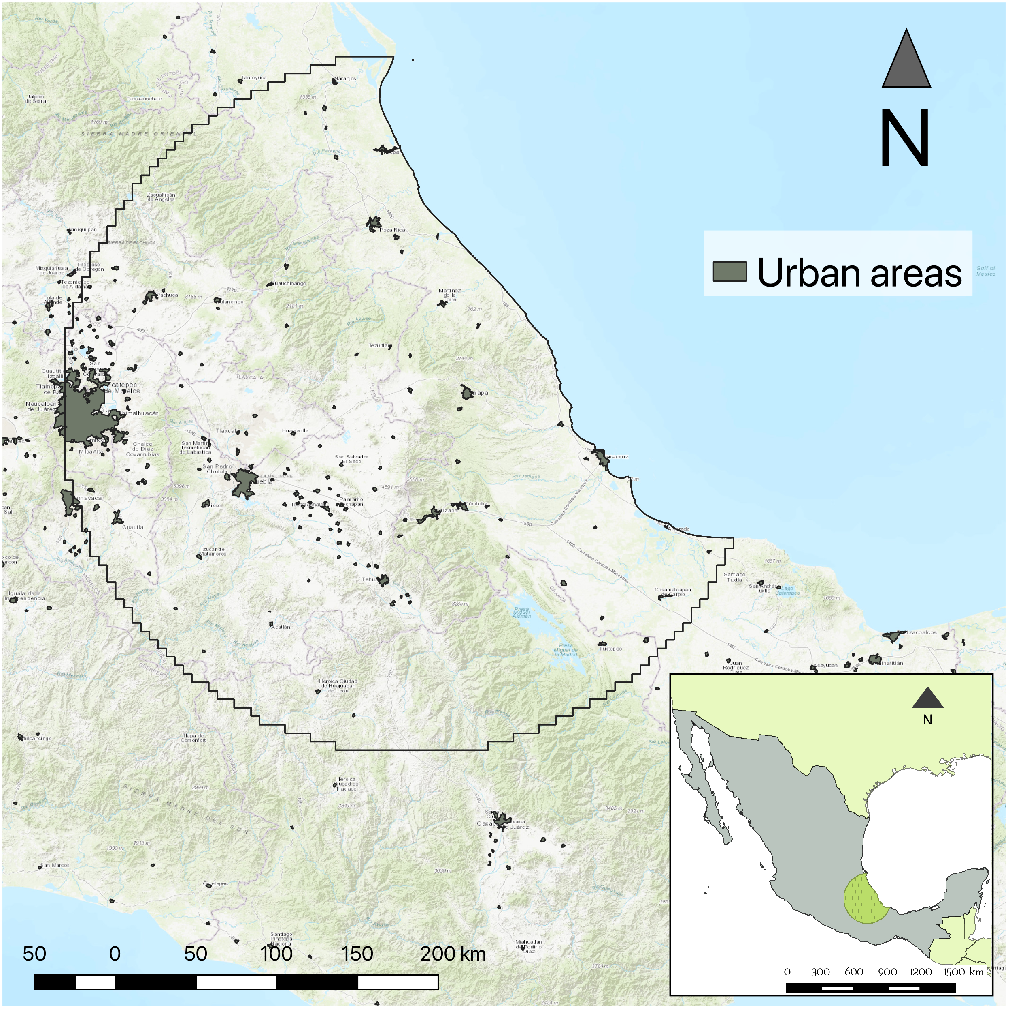
A map showing the study area (overlaid semicircular polygon) over central Mexico. Important cities are shown as grey polygons scattered across the area. Greener areas represent higher vegetation cover. The basemap used as background was obtained from the ESRI topographic tiling service.

### 3.2. Occurrence data

For the presence-only data we used the available GBIF occurrence data (GBIF Secretariat, 2015) registered before January 2015, constrained to the region 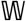. The raw data was downloaded from the GBIF portal with the catalog id: DOI:10.15468/dl.oflvla. Upon downloading, we performed a minimal data cleansing to remove records with missing information in any of the seven taxonomic ranks (i.e. kingdom, phylum, class, order, family, genus and species), acquisition date and collection code. We kept occurrences with identical coordinates as, historically, these occurrences might represent distinct different records collected in a common study area. Further information of this dataset, including all data attributions can be found in (GBIF.org, 2016).

We aggregated the occurrence data following the *choosing principle* described in subsection 2.2 to obtain response variables 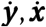 according to each example. The aggregation was by the class *Pinopsida* and kingdom *Plantae*, in the *Pines* example and, by the family *Tyrannidae* and class *Birds* for the *Tyrannids* case. Both examples used all known living records (*Life*) as background signal 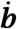. The taxonomic classification structure used was the GBIF Taxonomic Backbone (GBIF Secretariat, 2017).

### 3.3. Treatments for missing data

To assess the impact of using missing information in the prediction accuracy of the framework, we established two different treatments for fitting each model on each example. Recalling that both response vectors *Y* and *X* have entries of presence, relative absence and missing data, we defined the following treatments:

- treatment *i*: response vectors for the ToI (*Y*) and the sample (*X*) have missing data (i.e. 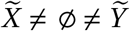).
- treatment *ii*: only the sample response vector (*X*) has missing data. That is, 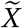 is the only source of missing information.

The motivation of using treatments is that they can serve as a middle hypothesis to assess the performance of the framework under scenarios with different proportions of missing data. The recommended scenario for use in practical applications is to use treatment *i*. We used the ROC-AUC estimate to measure the model’s performance within treatments. Using this estimate as an absolute measure between models may lead to wrong conclusions. For example, treatment *ii* implies that all the absences of *Y* are real and the sample *X* provides no information in the data augmentation methodology and therefore resulted in lower variance. This may lead to the conclusion that treatment ii performed better, and has greater predictive accuracy than treatment i. This conclusion would be true only under the assumption that the absences of the sampling effort are in fact true absences, which, in the case of presence-only data is false. Therefore, the comparison of presence-only models using the AUC-ROC estimate is only valid as a relative measure within models that used the same data, as it penalises models that estimate potential distributions (e.g treating absences as missing information) whilst favouring those that model realised distributions those where absences are informative) (Jiménez-Valverde, 2012). Comparing the AUC makes sense only when they are conditioned to a specific treatment and not between treatments.

### 3.4. Explanatory variables

The elevation data used were obtained from the Global Relief Model *ETOPO1* at 1 arc-minute resolution (Amante and Eakins, 2009). The precipitation data were obtained from the World Climatic Data *WorldClim* version 2 (Fick and Hijmans, 2017). The original data are composed in a raster model with c.a 1 km spatial resolution averaged from the years 1970 to 2000. The raster data were aggregated (by mean) to a scalar value for each areal unit in the spatial lattice equivalent to a spatial resolution of 4 km. This approach was used for the raster data. The distance to road dataset was generated in two steps. First we rasterised the National Road Network for Mexico (*Red Nacional de Caminos* (RNC) INEGI, Instituto Mexicano del Transporte and Gobierno de Mexico (2014), scale: 1: 250000) at 1 km spatial resolution. Later, we used this raster dataset to calculate its proximity to the closest road (pixels flaged as road) using the function `gdal_proximity` delivered as a stan-dalone command-line utility from (GDAL/OGR Contributors, 2018). The road network data were obtained from: Vázquez (2018). The population dataset was obtained from the WorldPop project (Sorichetta et al., 2015) for the year 2010. The dataset consists of population counts on each areal unit, each with a spatial resolution of 3 arc-seconds (c.a 100 m).

### 3.5. Data preprocessing

The occurrences, scenopoetic and anthropological data were spatially overlaid and aggregated on each areal unit of 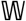. The aggregation method differed according to the data type. Mean and standard deviation were used for continuous variables, mode for categorical variables and the logical AND for binary data (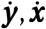 and 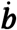). The data pipeline for processing the data was undertaken with *Biospytial* (Escamilla Molgora et al., 2020) a geospatial knowledge engine for processing environmental data https://github.com/molgor/biospytial.

### 3.6. Inference and prediction

We used a customised version of the R package *CarBayes* (Lee, 2013) and adapted it to fit models I, II and III. It includes a wrapper for easily fitting SDMs using one of the three models proposed using any type of fixed effects. The code is available from: https://github.com/molgor/CARBayeSDM. The package fits the model with a Markov Chain Monte Carlo (MCMC) method using a combination of Gibbs sampling and the Metropolis-adjusted Langevin Method (MALA), (Roberts and Tweedie, 2006). The posterior distributions were sampled by running 10000 iterations (using 5000 for burn-in) and a thinning interval of 5. Prediction for sites with missing information was done by sampling the posterior distributions of 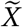 and 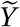. This same configuration was used in models I, II and III.

### 3.7. Comparison between models

Models I, II and III were compared with the *Deviance Information Criterion* (DIC) (Spiegelhalter et al., 2002). The DIC accounts for the number of parameters used and the likelihood of the observed data, given the statistical model assumed to be generating the data. The DIC is a generalisation of the Akaike information criterion (AIC) for hierarchical models, both measure the quality of the models in terms of their accuracy and parsimony. The DIC also serves as a Bayesian-based model selection tool. Model *A* is preferred to model *B* if its DIC value is lower than the one for *B* (i.e DIC_*A*_ < DIC_*B*_).

### 3.8. Comparison against Maxent

As mentioned in the introduction, we used the maximum entropy (MaxEnt) algorithm (Phillips et al., 2006) as a benchmark to compare the prediction accuracy of the proposed models. Contrary to models I, II and III, MaxEnt does not have a hierarchical specification and, therefore, calculating a DIC for model comparison is not possible. To address this limitation, we used a *k-fold* (*k* = 7) cross-validation methodology for measuring the quality of the predictions of all models. That is, on each fold, 1/7-th of the data was excluded from the fitting process and used as testing data to be compared against the corresponding predictions. This procedure was performed seven times, until every observation had a corresponding predicted value. We then used the *receiver operator characteristic* (ROC) curve and its area under the curve (AUC) (Fielding and Bell, 1997) as a measure of prediction accuracy. The same seven-fold cross validation was performed for models I, II and III with the difference that the excluded data were treated as missing data. The ROC / AUC values, as well as their corresponding 95% confidence intervals were calculated with the R package pROC (Turck et al., 2011).

Recalling that the proposed models are based on a spatial lattice structure (i.e. a CAR-based model), the spatial variation is modelled on a finite set of areal units. In the following case studies, these units were defined as square cells on a regular grid of approximately 4 km of spatial resolution. To make a fair comparison, we used the same spatial resolution and environmental values for fitting the MaxEnt models. Additionally, the background data (i.e. *pseudo-absences* in the MaxEnt jargon) used for fitting MaxEnt were obtained from locations with sampling observations but with no record of the taxon of interest, similarly to the sample selection bias for background data proposed by (Phillips et al., 2009). In other words, the *choosing principle* was also applied to the MaxEnt models resulting in the same input for all models (only valid for component *Y* (presence) of models I, II and III).

#### 3.8.1. MaxEnt optimisation

MaxEnt allows different configurations for model fitting. The most important are: the regularisation factor (reg) and the composition of mathematical transformations of the covariates, so-called *features* (see: Merow et al. (2013)). These features are equivalent to functions of the trend (i.e. they modify the fixed effect). To optimise the predictions of MaxEnt, we ran the 7-fold cross validation using different combinations of regularisation factors (reg ∈ (0.1,150)) and feature functions. In the case of the features, we used single and paired combinations of each of the following types: linear (l), quadratic (q), product(p), threshold (t) and hinge (h). The total number of different combinations (i.e models) for MaxEnt was 2250. The model was fitted with the R package maxnet (Phillips et al., 2017).

## 4. Results

### 4.1. Presence of Pines

We performed the methods described in section 2.2 to obtain response variables for Pines (*Pines*) and the botanical sample (*Plants*) using a geographical lattice 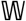 composed of 4060 cells (or unit areas). For the presence observations, 341 (8.4%) cells have known occurrences (class *Pinopsida*), 2559 (63%) have relative absences and 1160 (28.6%) are unknown (locations with missing observations). For the sample observations (botanical records), 2900 (71.4%) cells have known occurrence, 430 (8.4%) have relative absence and 730 (18%) unknown information (missing data).

The optimal MaxEnt, in terms of its higher predictive accuracy measured by the AUC-ROC was the one with a hinge feature type (nknots=50) and regularisation factor of 0.5. This combination, however, achieved the lowest predictions AUC of 0.67 ±(0.64,0.7)95% confidence interval (CI), when compared with models I, II and III (see figure 4a). Results from the best MaxEnt model and Models I, II and III are described in table 2.

**Table 1:**
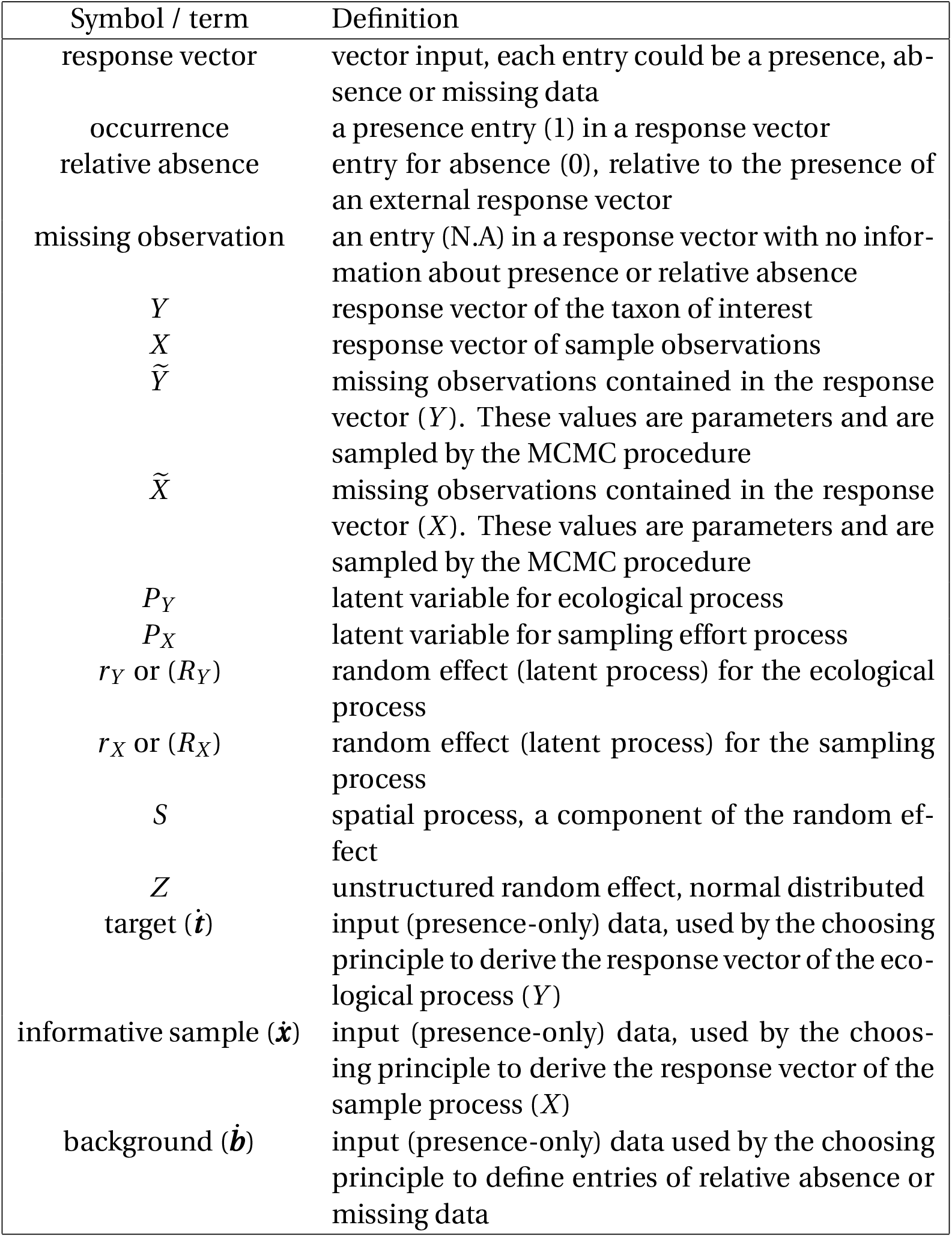
Definitions of the used terms and symbols

**Table 2:** Comparison of the presence-only models: Independent Spatial Components (Model 1), Common Spatial Component (Model 2), Correlated Spatial Components (Model 3) and Maximum Entropy (MaxEnt) for the presence of Pines (class *Pinopsida*) using botanical records (kingdom: *Plantae*) as sample effort. A 7-fold cross validation was performed to calculate the area under the receiver-operating characteristic curve (ROC-AUC) as a measure of quality for each model. Models with the * symbol were fitted using only missing data from *X* (sample), i.e. treatment *ii*.

| | DIC | ROC-AUC | 95% C.I | DIC$\star$ | ROC-AUC$\star$ | 95% C.I$\star$ |
| --- | --- | --- | --- | --- | --- | --- |
| Model I | 3517.6 | 0.835 | [0.81, 0.86] | **3421.2** | 0.874 | [0.85,0.89] |
| Model II | 3665.9 | 0.826 | [0.8,0.85] | 3647.9 | 0.877 | [ 0.86, 0.89] |
| Model III | **3440.2** | 0.832 | [0.80,0.85] | 3505.9 | 0.876 | [0.86,0.89] |
| MaxEnt | – | – | – | – | 0.67 | [0.64,0.7] |

For the treatment *i* (i.e. with both sources of missing information, see section 3.3), Model III (the one with correlated spatial structures) resulted to be the best ranked, that is, it achieved the lowest *Deviance Information Criterion* (DIC of 3440.2, see table 2). The predictive accuracy of this model, measured as the area under the ROC curve (i.e. AUC-ROC) was the highest of all three models (see figure 4a). The AUC of the three models fell within a common 95% credible interval of [0.8,0.86], that is, the predictive accuracy of models I, II and III was not significantly different.

Treatment *ii* (i.e. the one with no missing data in the sample effort component) produced slightly different results. In this case, Model I (independent spatial effects) was the best ranked by achieving the lowest DIC value (3421.2). The AUC in all models was higher than those on treatment *i*. However, in a similar way all of these values fell within a common 95% credible interval of [0.85, 0.89] (see supplementary materials fig: B.11). Possible reasons for this effect are explained in the next section. Additionally, the ROC curves in all models show similar variance described as the envelope of the ROC curve. Figures of this has been left to the supplementary materials (fig: B.11). The framework allows testing the significance the model’s parameters, in the same form as a Bayesian linear regression. In this sense, the variable *distance to road* was found to be the only significant covariate common to models I, II and III. That is, the zero is out of the 95% credible intervals (CI) of its posterior distribution. The scenopoetic variables (elevation and precipitation) were only significant in Model II. The selection of these specific covariates was based solely to demonstrate the capabilities of the model. As such, other covariates with stronger significance may be used further applications.

#### 4.1.1. Spatial results

Figure 3 shows the mean predicted latent surfaces for the presence of Pines *P_Y_* and sampling effort *P_X_* in all three models (left and right columns resp.). *P_X_* shows higher probability of occurrence than *P_Y_* across all the region. This is consistent in the three models. In contrast, the presence *P_Y_* revealed clustered patterns of high probability (figure 3). Of particular interest is the central zone that shows a high probability of occurrence. This area corresponds to the contact between the Eastern Sierra Madre and the Volcanic Axis and is of high elevation and high precipitation. In contrast, the MaxEnt model (fig: 3, bottom left panel) produced a smoother surface. The orographic features are more defined and the clustered patterns for presence are lost. Visual comparison between the models is difficult because of their similarity. However, in treatment ii (only one source of missing observations), Model II shows the compromise of estimating the sample *P_X_* to satisfy a common spatial component with *P_Y_*. In Model III, the median correlation obtained from the cross variance (Σ), between the presence of pines (*P_Y_*) and the sampling effort (*P_X_*), was 0.97 with (0.9,0.99) 95% credible interval. This result is consistent with the fact that the taxon of interest (i.e. pines) is totally contained in the sampling effort (i.e. plants). The complete estimates summary can be checked in supplementary section Appendix B.

**Figure 3:**
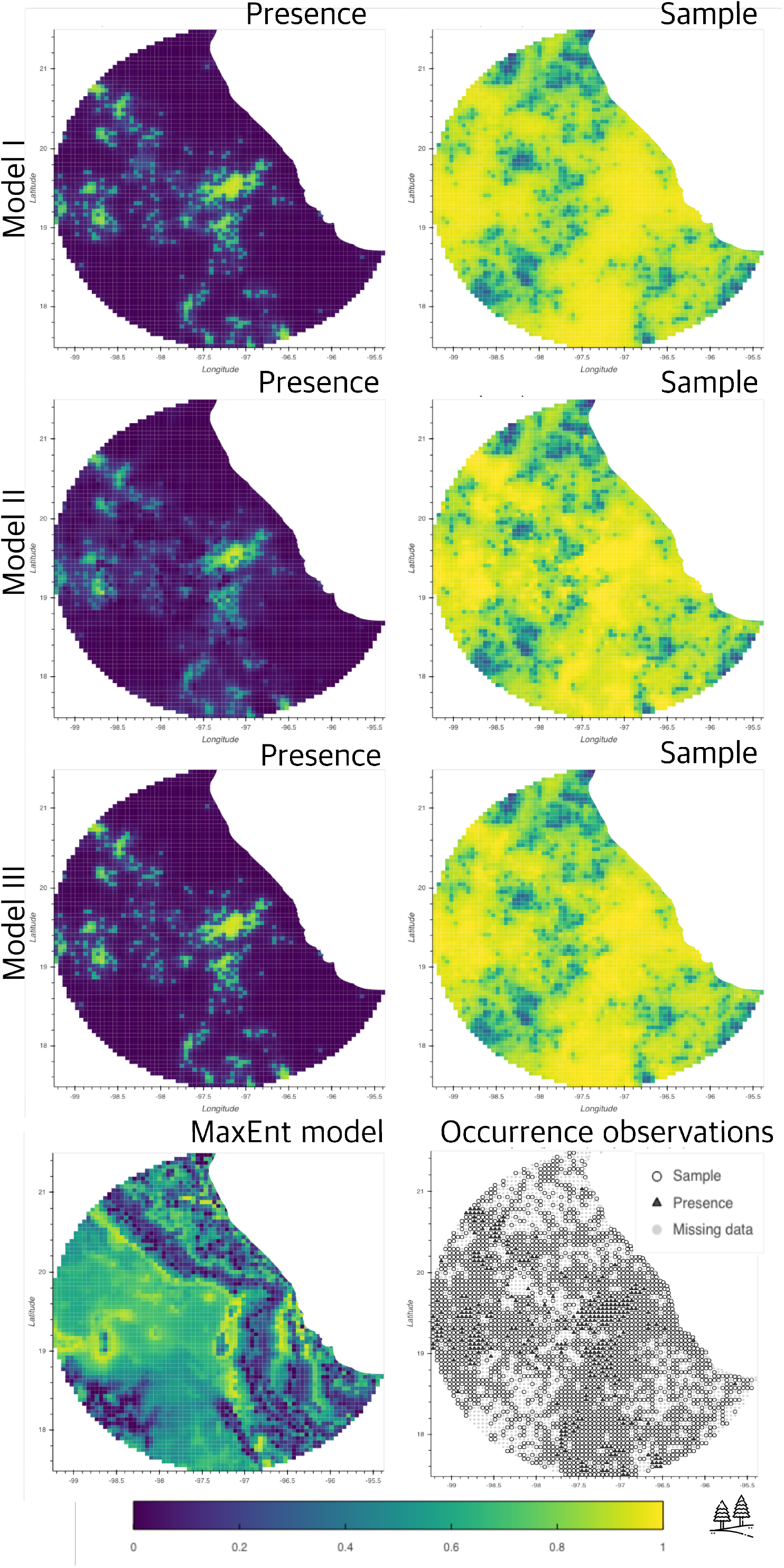
Comparison of models I, II and III against the maximum entropy algorithm (bottom left panel). The maps displayed here corresponds to the posterior mean probability for the three models using observations of pines as presence (panels on left) and botanical records (panels on right) as the sampling process. The bottom right panel shows the observations used to fit the models.

### 4.2. Results for the Presence of Flycatchers (family Tyrannidae)

This example was performed in the same study region (i.e., across the lattice 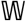). However, the data availability was significantly different and, therefore, the results were also different. In this example we obtained 596 (14.6%) cells with known occurrences of flycatchers, 368 (9.1%) with relative absences and 3096 (76.2%) of unknown or missing information. The occurrences for the sample (birds in general) was composed of: 990 (24.4%) known occurrences, 2340 (57.6%) relative absences and 730 (18%) missing data.

The optimal MaxEnt, in terms of its higher predictive accuracy measured by the AUC-ROC was the one with a combination of feature type of linear and threshold (nknots=50), and a regularisation factor of 0.7. The resulting optimal combination achieved a ROC-AUC of 0.61 ±(0.59,0.63)95% confidence interval (CI). The optimal parameter combination resulted to be equivalent to models I and III in terms of its predictive accuracy. That is, all the MaxEnt models are covered by the 95% confidence intervals of the ROC-AUC estimation for models I, II and III. Nevertheless, Model II (the one with a common spatial random effect) resulted to be significantly more accurate than the rest of the models. Figure 4b shows a comprehensive view of the aforementioned results. Additionally, a quantitative summary of these results is described in table 3.

**Figure 4:**
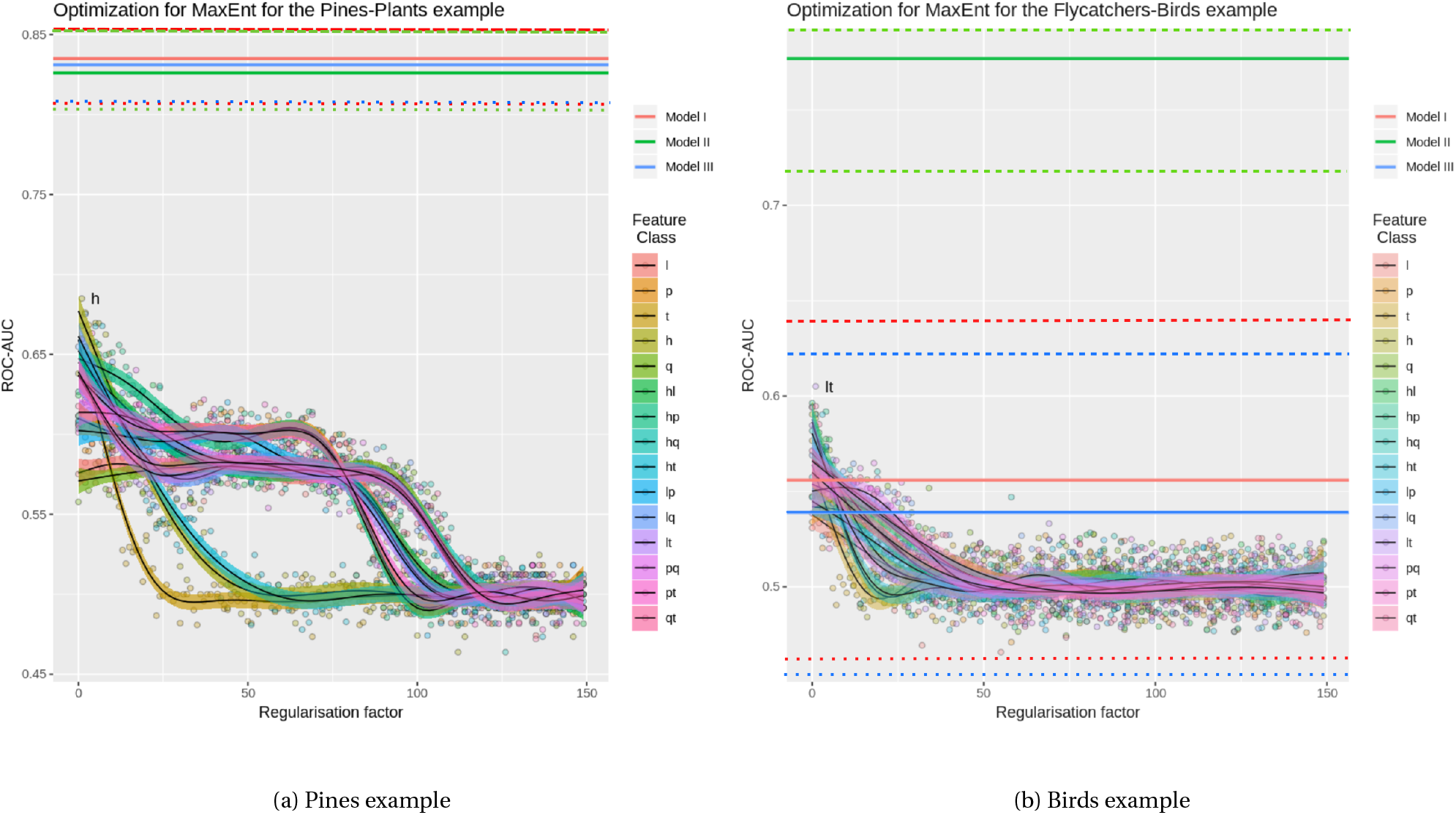
Area under the receiver operating characteristic curve (AUC-ROC) for the different models of the pines example (left panel) and the birds example (right panel). The dots in colours represent a MaxEnt models using different parameters of regularisation (x-axis) and feature type (vertical legend). The values in the y-axis correspond to the resulting AUC-ROC value according to that specific pair of parameters. The AUC-ROC values of models I (red), II (green) and III (blue) are shown as horizontal lines. Solid lines represent the mean AUC-ROC values for models I, II and III, while dotted and dashed lines represent their respective lower and upper (95%) confidence intervals.

**Table 3:**
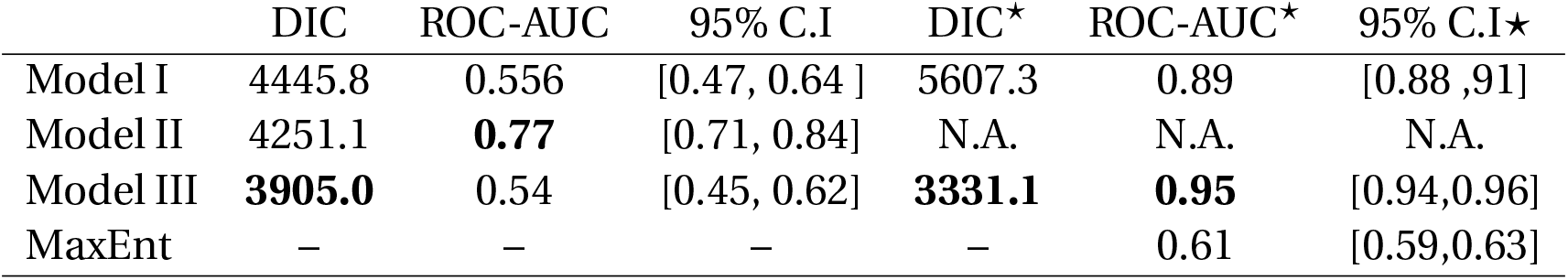
Comparison of the presence-only models: Independent Spatial Components (Model 1), Common Spatial Component (Model 2), Correlated Spatial Components (Model 3) and Maximum Entropy (MaxEnt) for the presence of the family *Tyrannidae* using birds as sample (class: *Aves*). A 7-fold cross validation was performed to calculate the area under the receiver-operating characteristic curve (ROC-AUC) as a measure of quality for each model. Models with the * symbol were fitted using only missing data from *X* (sample), i.e. treatment *ii*.

In treatment *i* (i.e. missing data in both response vectors, the one for presence and the one for sample), Model III (correlated spatial components between the ecological process and the sampling effort) was the best ranked, achieving the lowest DIC value (3905), similarly to the Pines example. However, its accuracy in terms of ROC-AUC was close to random classification, reaching an AUC of 0.54 with ± (0.45,0.62) at 95% CI. Model I (independent spatial effect for the ecological and the sampling components) obtained similar values of ROC-AUC (0.56 ± (0.47,0.64) at 95% CI). In contrast, Model II obtained the highest predictive accuracy ( 0.77 ± (0.71,0.84)) with a DIC of 3905, second in rank. (see figure 4b); In addition, models I and III achieved a low predictive power compared to the benchmark model (MaxEnt).

Treatment *ii*, (i.e only one response vector (*X*) with missing information) showed contrasting results. Although model III (correlated components) ranked best, in terms of a lowest DIC (3331.1), its AUC was 0.95 ± (0.94,0.96). Model I (independent spatial components) followed with an AUC of 0.89 ± (0.88,0.91). Model II, could not obtain valid posterior distributions, as its log-likelihood diverged to -∞. We discuss possible reasons and circumventing strategies in the next section.

All results are shown in table 3. Based solely on the DIC, Model III was ranked first in both treatments. However, in cases with large proportions of missing data (as in treatment *i* with 76.2% cells) the prediction accuracy (ROC-AUC) was low. This effect highlights the importance of selecting informative missing data as well as the type of model to use. These issues are explored further in the discussion section.

The covariate *Distance to roads* was found to be significant in models I and III. The rest (elevation, precipitation and population count) were not significant in all three models. The selection of these specific covariates was based solely to demonstrate the capabilities of the model. As such, other covariates with stronger significance may be used.

#### 4.2.1. Spatial results

Figure 5 shows the mean predicted latent surfaces for the presence of flycatchers *P_Y_* (*Tyranids*) and relative sample *P_X_* (*Birds*) in all the three models (left and right columns resp.). Model I presents a clear difference between *P_Y_* and *P_X_* (figure 5, first row). In this case, *P_Y_* appears more smooth with patches of lower probability, although always with probability higher than 0.2. The surface *P_X_* in model I (fig: 5, top right panel) has clear shaped patterns with contrasting probabilities between interior regions (*pocket shapes*). This feature is present in both surfaces of model II (fig:5, second row) and model III (fig:5, third row) The fixed effects (covariates) for *P_X_* and *P_Y_* are close to zero, therefore, the spatial variation is driven only by the common structure *S*. In the case of model III, the sample surface *P_X_* presents greater connectivity and higher probabilities in places with known observations. Both surfaces, however, present a similar structure in shapes and patterns.

**Figure 5:**
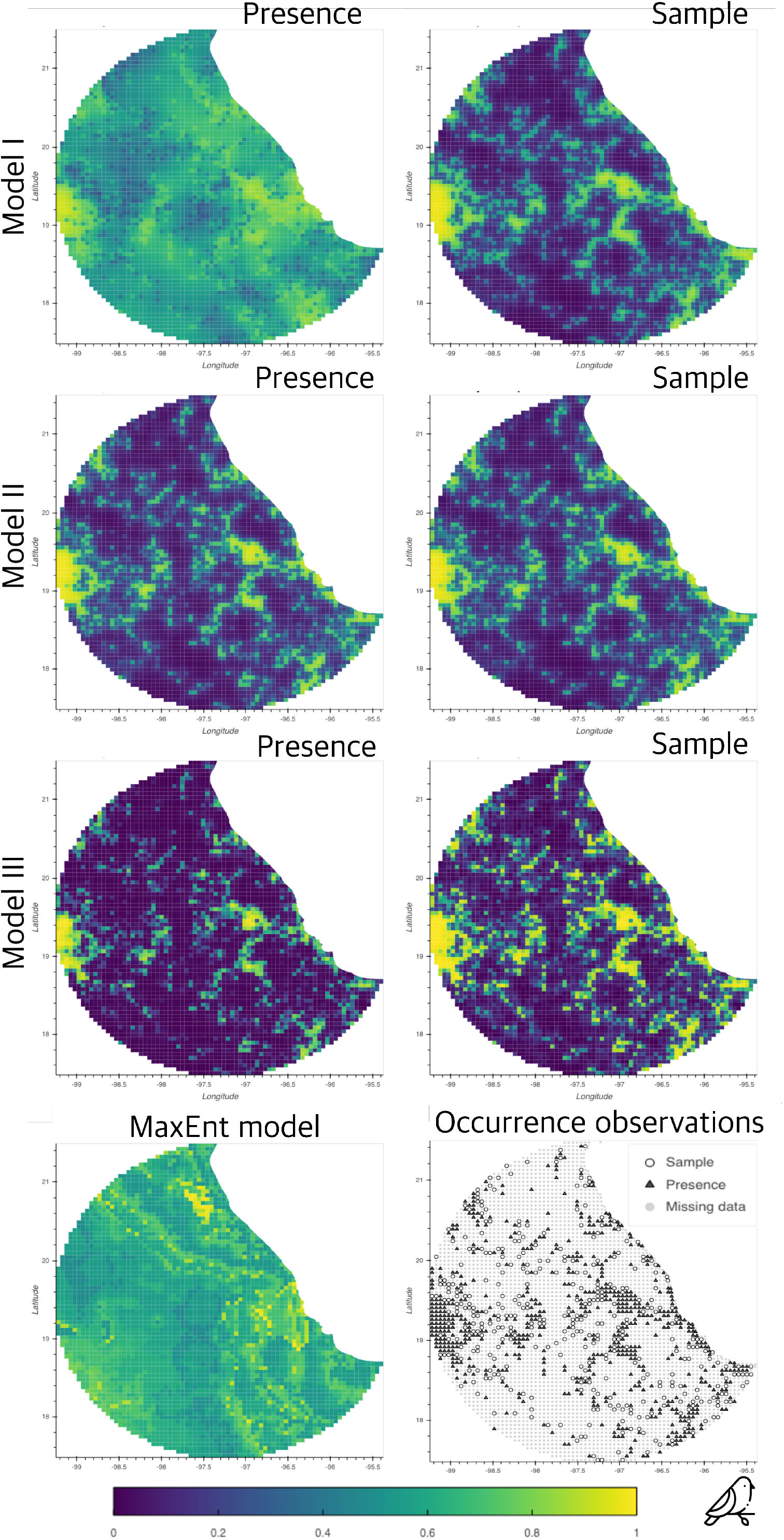
Comparison of models I, II and III against the maximum entropy algorithm (bottom left panel). The maps displayed here corresponds to the posterior mean probability for the three models using observations of flycatchers as presence (panels on left) and observations of birds records (panels on right) as the sampling process. The bottom right panel shows the observations used to fit the models.

In contrast, the MaxEnt prediction lacks the random spatial effect component. The resulting probability surface is determined exclusively by the features used by the covariates. Although is possible to distinguish spatial patterns within the region, the predicted probability is in general close to uniform random classification (i.e. 0.5). This effect is supported by the obtained AUC-ROC value of the cross-validation analysis (0.6) (fig: 4b (a)). In Model III, the median correlation, obtained from the cross variance (Σ) between the presence of flycatchers (*P_Y_*) and the sampling effort (*P_X_*), was 0.996 with (0.993,0.998) 95% credible interval. As in the latter example, this result is consistent with the fact that the taxon of interest (i.e. flycatchers) is totally contained in the sampling effort (i.e.birds). The complete estimates’ summary can be checked in Appendix C.

## 5. Discussion

The bivariate CAR modelling framework uses an additional source of information, apart from the presences of the target species. This extra information comes from sampling observations related to other species and other taxa that, according to the modeller, give complementary information relative to the occurrence of the taxon of interest (ToI). The framework relies on three fundamental concepts: *i*) the sampling effort as complementary information for inferring the probability of presence, *ii*) the spatial autocorrelation structure for determining the variability and occurrences likelihood across the landscape, and *iii*) the *choosing principle*, a mechanism for determining presences, relative absences and missing data from presence-only records. Both examples showed that, at least one of the three proposed models outperformed MaxEnt. The results in tables 2 and 3 show that the models’ goodness-of-fit statistic (i.e. DIC) and predictive accuracy increased in treatment *ii*, that is, when the absence of records were treated as real absences. This is expected because assuming missing data as real absences reduces uncertainty.

These results show that the proportion of missing data plays a fundamental role in the predictive capability of the model. This effect is recognised in the flycatchers example, where the proportion of missing observations is much higher (76% of the total number of regions) compared to presences and relative absences. In this case, models I and III produced low predictive accuracy, similarly to MaxEnt, with an AUC-ROC of near 0.6 (i.e., close to random classification). In contrast, model II, although ranked second in terms of DIC, achieved the highest predictive accuracy (AUC-ROC). This result is also supported by by the high number of missing data (increased uncertainty) and reduced number of spatial parameters to fit. In terms of models’ parsimony, one shared spatial latent effect (model II) has less parameters to fit compared with two spatial effects in the case of models I and II.

The three proposed models impose different restrictions on how the spatial autocorrelation structure affects the probability of a species to occur. The more complex the spatial structure is, the more presence-only observations (and less missing data) are needed. This can be modulated by the amount of missing data with respect to the relative absences determined by the sampling effort observations and the choosing principle. Consequently, using an appropriate informative sample becomes crucial for obtaining accurate inferences and predictions. This finding highlights interesting paths for future research: one related to the selection of informative observations for the sampling effort process, and the other for different choosing principles.

Model II may be a better alternative for taxa with sparse spatial distributions and large proportion of missing data. Nevertheless, model II presented problems with identifiability in treatment *ii* (i.e. missing data only in the ToI observations and assumed real absences in the sampling process). A possible reason is that the inference method could not find a suitable compromise in accounting for a common spatial effect that had two constraints. One, the accountability of residuals of both processes (*P_Y_* and *P_X_*) and two, the restrictions imposed by the intrinsic CAR model specification. That is, the sum of the random effect on all the lattice areas should sum one. A possibility to circumvent this last restriction is to specify, instead, a proper CAR model (e.g (Leroux et al., 2000)). The package CARBayes (Lee, 2013) allows this specification. We recommend the practitioner to compare the three models accordingly to fit specific needs.

### 5.1. The role of the choosing principle

When presence-only data are used, any choosing principle is inevitably a source of potential bias. Thus, the research question and the selection of the sampling effort observations play a fundamental role in determining the accuracy of predictions. The way relative absences and missing data are derived implies ecological assumptions that should be kept in mind when one tries to model species (taxon) distributions. For example, following the *biotic, abiotic, movements* (BAM) diagram proposed (Soberon and Nakamura, 2009), if the objective is to model the *realised distribution*, (i.e., places where the species lives in reality) absences become informative. If on the other hand, the objective is to model the species’ *potential distribution* (i.e. places where it can survive and thrive due to suitable environmental conditions) absences may constitute missing data. See equivalent concepts from a SDM approach Jiménez-Valverde et al. (2008).

In our framework, we used the sample observations *X* together with the *choosing principle* to discriminate between informative absences and missing data. If the sampling effort is chosen to be informative it can increase significantly the accuracy of predictions (see table 2).

The current choosing principle assumes that for every location *k*, if the ToI (e.g. species) is not present, but the sample observation exists (*X_k_* = 1), then the ToI is assumed to be absent (*Y_k_* =0). In some applications this assertion may be incorrect and, if the sample observations *X* consist as well of presence-only data, the bias in false absences can propagate in both processes. This problem is present in all presence-only methods that tries to account for the sampling bias using pseudo-absences (e.g. target-background approach of Phillips et al. (2009)), given the intrinsic bias of the collected data. Ideally, the best way to rank distinct choosing principles, given a ToI, is using presence-absence data. The proposed choosing principle is not intended to be a general rule for all species and problems. An it is worth for the modeller to consider other choosing principle in which relative absences and missing data can be specified from presence-only data. For example, another type of choosing principle can incorporate information on other species features. For example movement, since the accessibility of an area can be indicative of poor sampling and its use has been shown to reduce bias in occurrence data (Monsarrat et al., 2018).

We would like also to explore further the role of the taxonomic structure in determining informative samples. In the examples we used broad and generic groups, jumping from class *Pinopsida* to kingdom *Plantae*, in the case of Pines, and from family *Tyranidae* to class *Aves*, in the case of the flycatchers. We hypothesise that using the immediate parent node of the ToI, according to its taxonomical classification, could give more accurate models for certain groups. An example of this could be the use of the family (of the ToI) as sample, if the ToI is a type of genus.

In recent years, spatial point process (SPP) models have been proposed to model presence-only occurrences (see Velázquez et al. (2016) for review).

This is a sensible choice of modelling giving that these models are able to represent discrete events in a continuous space. Recently, authors like (Renner et al., 2015, 2019) proposed a combined likelihood approach for modelling the spatial dependence using a latent log Gaussian Cox process (Møller et al., 1998). Although these models are sound and have been used satisfactory, the assumptions about the required sample design restrict their application to only specific cases (Gelfand et al. (2013), Chp. 20). Additionally, in SPP models, all information is contained in the location of the occurrences and separating the sampling effort from the ecological process, can lead to confounding and identifiability problems. In our opinion the use of spatial lattices (i.e. Gaussian Markov random fields) for modelling spatial autocorrelation presents a more appropriate alternative for modelling generic species.

### 5.2. Advantages in using this framework

The model is defined in a spatial lattice. The observations occurred on a given area element can be aggregated to reflect presences or abundances. That is, the model support repeated measurements within areas. In addition, the probabilities for presence in areas that have not been sampled can be inferred by the neighbouring areas. The method is able to infer places where data availability is limited. The model specifies a Bayesian hierarchical model and accounting uncertainties of the parameters is possible. This brings the possibility to perform hypotheses testing on the posterior sample. As it is a hierarchical model it is possible to perform model selection using the DIC statistic. The structural components of the models, that is, the ecological process and the sampling effort can be explicitly modelled using different covariates and even feature classes, as the ones used by MaxEnt. Lastly, the choosing principle provides a flexible form to assign absences and missing data.

### 5.3. Limitations

Manipulating the spatial random component of the model implies greater computational complexity on the order of *O*(*n*^3^) (in its worse scenario). Although, the matrix is sparse and the inference uses optimised numerical methods that can reduce the computational complexity, the numerical methods involved are more intensive than MaxEnt or other models that are not based on hierarchical Bayesian inference. This is a limitation for studies that requires extended regions involving hundreds of thousands of area elements.

Another limitation is that the specification of the spatial effect is based on discrete spatial distributions. This implies that, once the model is fitted, it is not possible to make predictions on observed regions or data (as opposed to geostatistical models). Also, depending on the specification, a modeler may need the spatial random effect to be continuous in space, instead of over a discrete lattice. If this is the case we recommend the use of SPP-based models like (Renner et al., 2015, 2019).

### 6. Significance Statement

The presented work provided three alternatives to model the spatial distribution of species using solely observations of presences. The two case studies showed that, in terms of predictive accuracy, at least one of the alternatives outperformed the most popular method for modelling species distributions (i.e MaxEnt).

The framework can be applied in a variety of problems where information on species absences is unknown but data from other species is available. As this approach returns posterior probability distributions, it provides valuable information for performing spatial analyses, estimating predictions and uncertainties and testing hypotheses related to the model’s parameters.

## 7. Data and source code availability

Currently the code and data are stored in the following repository: https://github.com/molgor/CARBayeSDM. We intend to put the code and data in a long term curated repository such as Dryad or FigShare.

## Acknowledgments

This project was jointly sponsored by the Mexican Science and Technology Council (CONACyT) under the doctoral program: *Becas al Extranjero* and the Faculty of Science and Technology from Lancaster University. Icons for birds and pines made by Freepik from www.flaticon.com.

## 9. Biosketch

Juan Escamilla Mólgora is interested in developing computational and statistical methodologies for studying spatial patterns of life at different scales. This work was part of his PhD research at Lancaster University on the development of a computational and statistical framework for modelling species distributions using presence-only data from different sources. The co-authors collaborate in developing spatial statistical methods applied to epidemiological and environmental problems.

### 9.1. Authors’ contributions

All authors developed the general framework and provided critical feedback in all the stages of this work. More specifically, PD proposed the three model specifications. PA proposed the choosing principle. LS and JEM designed the modelling and simulations strategies. JEM prepared the data, implemented the models, performed the analysis and visualizations and wrote the manuscript with inputs and edits from all co-authors. PA, LS and PD supervised the project.

## Appendix A. Supplementary materials I: Framework specification

We begin by defining a grid inside a region of interest located somewhere on the Earth’s surface. Mathematically this is a spatial lattice 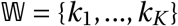 that partitions a compact set 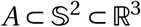 into *K* non-overlapping compact subregions. Let 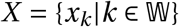 be the recorded presence of a certain sample (or survey) and 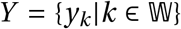 the presence of a taxon (e.g. species) of interest (ToI). As such, *x_k_* and *y_k_* are two binary random variables corresponding to the events of: *a sample x_k_ has been registered in location k* and *taxon y_k_ is present at location k*. Missing observations are defined in the same lattice as: 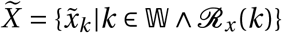 where 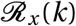 is the predicate of: *there is no recorded evidence of x in k* and similarly, 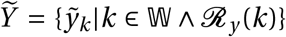 where 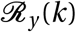 is the predicate of: *there is no recorded evidence of the presence of y in k*. The data augmentation methodology (Tanner and Wong, 1987) implemented in CARBayes (Lee, 2013) generates posterior samples of 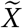 and 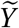. We opted to omit any further specification for the variables 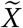 and 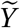 here, to simplify the description of the framework.

The general specification of the framework factorises the joint probability distribution in the following form:

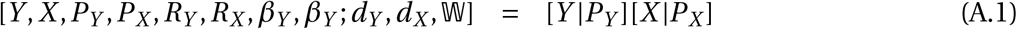

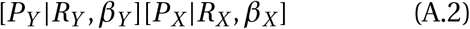

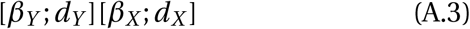

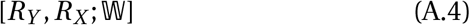

Equations 1 to 3 are consistent across the framework while the specification for equation 4 (i.e. *random effects*) vary according to three different assumptions of spatial autocorrelation; independent components (model I), a common spatial component (model II) and correlated spatial components (model III). We start by defining equations 1 and 2. That is, the probability of presence for a ToI (*Y_k_*) given the latent variable *P_Y_*(*k*) in a cell *k* and similarly, the probability of a sample *X_k_* to be present given its respective latent variable *P_X_*(*k*). These binary random variables are modelled as following:

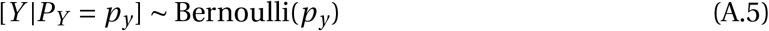

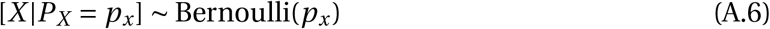

### Appendix A.1. Latent variables P_Y_ and P_X_

We assume that the presence-only data represent realizations of a joint stochastic process separable in two components: one relative to an ecological process *P_Y_* that drives the environmental suitability for the ToI, and another process *P_X_* related to the sampling effort. We, therefore, model [*P_Y_* = *p_y_*|*R_Y_* = *r_y_, β_Y_; d_Y_*] and [*P_X_* = *p_x_*|*R_X_* = *r_x_, β_X_; d_X_*] (eqs. A.2) according to the following specification:

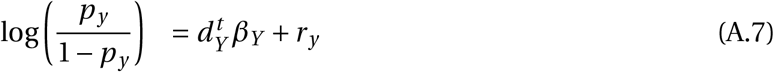

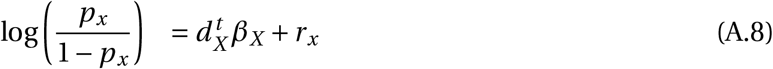

where *d_X_* and *d_Y_* represent vectors of explanatory variables and *r_X_* and *r_Y_* the random effects for *X* and *Y* respectively. Specifically, *d_Y_* is suited for environmental variables of ecological importance, while *d_X_* should account for variables that help explain the sampling process. The prior distributions for *β_Y_* and *β_X_* (eq: A.3) are defined, as default, as uninformative zero-mean normal distributions with default variance 100,000. We acknowledge that the use of uninformative priors can yield to skewed parameter estimates and negate the advantage of using Bayesian methods over frequentist analyses (Hobbs and Hooten, 2015; Gelman and Shalizi, 2013). These hyperparameter values are default options in CarBayes (Lee, 2013) and, consequently, in our modelling framework. As such, they can be changed according to the user needs. See (Lemoine, 2019) for a concise guide on using informative and weakly informative priors in ecological models. In the following section we present the three alternatives for modelling *R_X_* and *R_Y_*.

### Appendix A.2. Random effects

The general form of the random effects component for *P_Y_* (and *P_X_*) is defined as an independent zero-mean random variable *R_Y_* (*R_X_*). This variable accounts for the combined effect of a spatial process *S_Y_* (*S_X_*) that models the spatial variation across the lattice 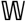 and an independent normally distributed random variable *Z_Y_* (*Z_X_*) with variance 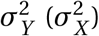 that accounts for unstructured noise inside each cell of the lattice.

Specifically, these random effects are defined as follows:

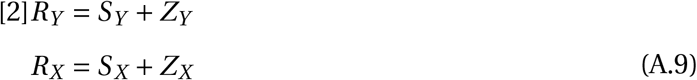

where *Z_Y_* ~ *N*(0, *σ_Y_*) and *Z_X_* ~ *N*(0, *σ_X_*) and the spatial components *S_Y_* and *S_X_* are modelled as *intrinsic conditional autoregressions* (ICAR) (Besag, 1974; Besag et al., 1991) with parameters 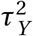 and 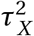 respectively, over the lattice 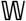. In the rest of this work we represent 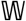 in its matrix form, that is, the adjacency matrix *W* of its graph representation; defined as a *k* × *k* symmetric matrix with entries: *w_i, j_* = 1 = *W_j,i_* if cells *i* and *j* are neighbours, otherwise *W_i, j_* = 0. Modelling the spatial autocorrelation as an ICAR eases significantly the computation of *W*^-1^ with the aid of optimised methods for sparse matrix algebra (Rue and Held, 2005). This approach simplifies significantly the inference, prediction and posterior sampling, a great advantage in applications with large datasets.

### Appendix A.3. Three models for spatial autocorrelation

The proposed framework assumes that the ecological process *P_Y_* and the anthropogenic sampling process *P_X_* are independent when conditioned to the random effects *R_Y_* and *R_X_* (see figure 1 and eq: A.2). This assumption implies that the only source of dependency between *R_Y_* and *R_X_* is the dependency between the spatial effects *S_Y_* and *S_X_*, this by the assumption of independence between variables *Z_Y_* and *Z_X_*. Moreover, the framework assumes that the observations of presence for the ToI and the existence of the survey (sampling) are independent when conditioned to the spatial effect. As such, the spatial autocorrelation structure is the component responsible for informing both processes. In order to test for this we designed three possible models in which the spatial processes *S_Y_* and *S_X_* inform *R_Y_* and *R_X_*. Model I in which the spatial components *S_Y_* and *S_X_* are independent, Model II with a unique spatial component shared between both processes *P_X_* and *P_Y_* (i.e. *S_X_* = *S_Y_*) and Model III in which the spatial components *S_X_* and *S_Y_* are correlated. Below we give the full description of each model.

#### Appendix A.3.1. Model I: Independent Spatial Components (ISC)

This model assumes that the spatial random effects on both processes (*R_X_, R_Y_*) are independent. By equations A.9 the joint distribution is given by

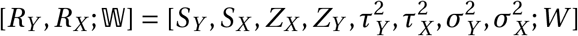

and, given the assumptions on independence, it can be factorised into:

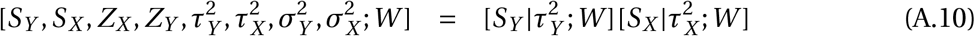

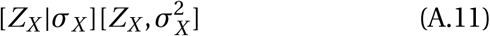

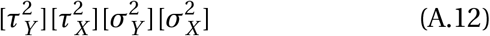

where the term 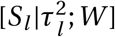 (*l* being *X* or *Y*) is modelled as an ICAR (Besag, 1974; Besag et al., 1991) with a full conditional form of:

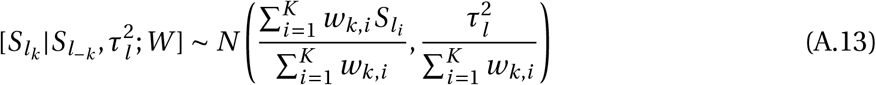

for each process *l* ∈ {*Y, X*} on each cell *k* (i.e. *S_lk_*). The prior distributions for parameters 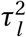 and 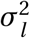 are defined as inverse gamma(1,0.01), default values in the package *CARBayes*. Figure 1a (in the main text) shows a general DAG structure for this model.

#### Appendix A.3.2. Model II: Common Spatial Component (CSC)

This model assumes that the random effects *R_X_* and *R_Y_* share the same spatial component *S* (i.e. *S_X_* = *S_Y_*). By equations A.9 the joint distribution is given by 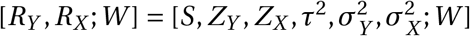 and, given the assumptions on independence, it can be factorised as:

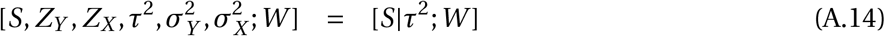

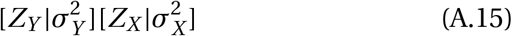

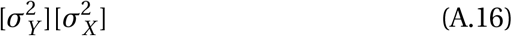

Similarly to model I, the spatial effect [*S*|*τ*^2^; *W*] is modelled as an ICAR (Besag, 1974; Besag et al., 1991) in full conditional form on each cell 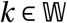.

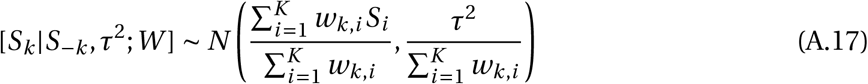

The prior distributions for parameters 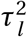 and 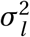 are defined as inverse gamma(1,0.01), default values in the package *CARBayes*. Figure 1b (in the main text) shows a general DAG structure for this model. Model II is specified as a two-level model where each areal unit *k* has two response variables, *X_k_* and *Y_k_*. The individual level variation is split into two groups: *Z_X_* and *Z_Y_*. Figure 1b shows the DAG describing the model.

#### Appendix A.3.3. Model III: Correlated Spatial Components (CSC)

This model specifies the joint random effect [*R_Y_, R_X_; W*] as a combined effect of the spatial processes, *S_Y_* and *S_X_*. To model this effect, both spatial effects are ensembled as a bivariate conditional autoregresive (BCAR) process that accounts for both *S_Y_* and *S_Y_* simultaneously. To improve the identifiability of the model, the unstructured random effect (i.e. *Z_X_* and *Z_Y_* in models I and II) is integrated into the spatial effect using a more relaxed specification of the spatial autocorrelation structure. This specification, proposed by Leroux et al. (2000), adds a new parameter *ρ* that models the strength of the spatial dependency. When *ρ* = 1 the spatial dependency is maximum and the spatial process is equivalent to an intrinsic CAR model. On the other hand, if *ρ* = 0 there is no evidence of spatial autocorrelation and therefore, the observations are spatially independent. To make the comparison between models I and II consistent, we have restricted *ρ* = 1. However, this restriction can be removed according to the needs of the users. Following the equations A.9 and the DAG specification shown in figure 1c (in the main text) the joint distribution [*R_Y_, R_X_*; *W*] can be factorised as:

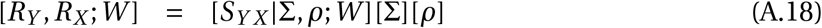

The combined random effect *S_YX_* is defined as the Kronecker product between the Leroux et al. (2000) CAR model and a 2 × 2 covariance matrix Σ that accounts for the cross variable effect between both processes. The correlation between both variables can be calculated as:

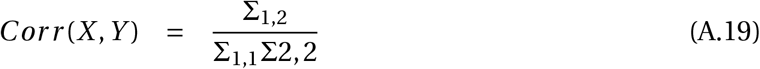

The BCAR model is a particular case of the multivariate model (MCAR) proposed by Gelfand and Vounatsou (2003) and it has been implemented in the R package CARBayes (Lee, 2013) following the proposal of Kavanagh et al. (2016). *S_YX_* is a realization of the following multivariate normal distribution:

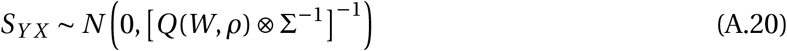

The autocorrelation function *Q*(*W, ρ*) is defined by the precision matrix:

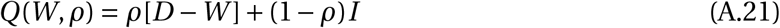

where *D* is a *k* × *k* diagonal matrix in which each entry *d_i,i_* is equal to the number of neighbours of each unit area *i* ∈ {1,‮‮,*k*}. The prior for Σ is distributed as Inverse-Wishart(3,Ω) with three degrees of freedom and Ω = *I*_2×2_ as scale matrix. The prior [*ρ*] is a non-informative uniform (0,1) distribution. The DAG describing the model is described in figure 1c.

## Appendix B. Supplementary materials II

This section contains the summary statistics of the fitted posterior distributions of the parameters corresponding to models I, II and III, described in summary in the main text (section: 2) and extensively in the supplementary materials Appendix A. The summary statistics corresponding to the presence of pines (using plants as sampling effort) is showed first. The second case study is showed in the next section. The structure of every table is the same for all models in both examples. The rows describe the parameters corresponding to each model (on each table). The first three columns describe the median, upper and lower bounds of the 95% credible intervals. The `n.effective` column indicates an estimate for the size of independent samples (taking into account autocorrelations within each chain of the MCMC sampler). The column % `accepted` refers to the proportion of times a proposed value was accepted by the Metropolis updating step as a new value of the posterior sample (see (Lee, 2013)). The column `Geweke.diag` refers to Geweke’s convergence diagnostic (Geweke, 1992) which compares the means calculated from distinct parts of the Markov chain to test for convergence of the stationary distribution (default first 10% and last 50%). If the chains reached a stationary distribution, then the two means are equal and Geweke’s statistic has an asymptotically standard normal distribution. All models can be fitted in CARBayes (Lee, 2013), which uses the R package Coda (Plummer et al., 2006) for calculating `n.effective` and `Geweke.diag`.

### Appendix B.1. Estimates for the predicted presence of Pines using botanical records as sample

**Table B.1:** Posterior summaries of all the parameters in Model I with the associated 95% credible intervals for the example of pines. Parameters 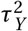 and 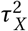 correspond to the variance of the spatial effects of the presence (Y) and the sample process (X) (i.e. *S_Y_* and *S_X_*) respectively. Likewise, 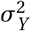 and 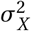 correspond to the variance of the unstructured processes *Z_Y_* and *Z_X_* respectively. Significant parameters are shown in **bold**. For further information see section: 3

|  | Median | 2.5% | 97.5% | n.sample | %accept | n.effective | Geweke.diag |
| --- | --- | --- | --- | --- | --- | --- | --- |
| (Intercept of Y) | -1.1871 | -4.0872 | 0.9928 | 10000 | 64.2 | 16.0 | -7.8 |
| Elevation | 0.0002 | -0.0002 | 0.0006 | 10000 | 64.2 | 299.9 | -2.0 |
| Precipitation | 0.0002 | -0.0001 | 0.0005 | 10000 | 64.2 | 206.4 | 0.4 |
| $\tau_Y^2$ | 19.6638 | 13.2754 | 45.1344 | 10000 | - | 8.5 | -1.3 |
| $\sigma_Y^2$ | 0.3658 | 0.0357 | 0.7923 | 10000 | - | 3.1 | 1.8 |
| **(Intercept of X)** | 3.0309 | 2.4178 | 3.9749 | 10000 | 61 | 24.3 | -0.9 |
| **Dist. to road** | -0.0002 | -0.0004 | -0.0001 | 10000 | 61 | 1294.1 | 0.5 |
| Population | 0.0000 | -0.0001 | 0.0001 | 10000 | 61 | 1320.2 | 0.4 |
| $\tau_X^2$ | 5.2708 | 2.7058 | 9.5806 | 10000 | - | 8.7 | -1.1 |
| $\sigma_X^2$ | 0.1818 | 0.0637 | 0.3250 | 10000 | - | 7.9 | -1.1 |

**Table B.2:** Posterior summaries of all the parameters in Model II with the associated 95% credible intervals for the example of pines. The parameter *τ*^2^ represents the variance of the common spatial effect. Parameters *σ*^2^ and *σ*^2^ correspond to the variance of the unstructured process *Z_Y_* and *Z_X_*. Significant parameters for the fixed effect are shown in **bold**. For further information see section: 3

|  | Median | 2.5% | 97.5% | n.sample | %accept | n.effective | Geweke.diag |
| --- | --- | --- | --- | --- | --- | --- | --- |
| **(Intercept)** | -0.7085 | -1.0766 | -0.3426 | 5000 | 51.6 | 80.5 | -4.9 |
| **Dist. to road** | -0.0002 | -0.0004 | -0.0001 | 5000 | 51.6 | 170.9 | -1.2 |
| Population | 0.0000 | -0.0001 | 0.0001 | 5000 | 51.6 | 150.2 | -0.2 |
| **Elevation** | 0.0002 | 0.0000 | 0.0004 | 5000 | 51.6 | 79.7 | 1.6 |
| **Precipitation** | 0.0003 | 0.0001 | 0.0004 | 5000 | 51.6 | 85.9 | 3.5 |
| $\tau^2$ | 6.8838 | 4.7169 | 11.8695 | 5000 | - | 5.5 | 5.1 |
| $\sigma^2$ | 9.7797 | 2.8682 | 72.7988 | 5000 | - | 5000.0 | 1.1 |

**Table B.3:** Posterior summaries of all the parameters in Model III with the associated 95% credible intervals for the example of pines. Parameters 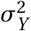 and 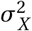 correspond to the variance for the presence (*Y*) and the sample (*X*). The term corr*_X,Y_* indicates the correlation between these two processes. Significant parameters for the fixed effect are shown in **bold**. For further information see section: 3

|  | Median | 2.5% | 97.5% | n.sample | %accept | n.effective | Geweke.diag |
| --- | --- | --- | --- | --- | --- | --- | --- |
| **(Intercept of $Y$)** | -7.7938 | -9.2851 | -6.3099 | 5000 | 55.6 | 60.5 | 6.4 |
| Elevation $Y$ | 0.0003 | -0.0001 | 0.0007 | 5000 | 55.6 | 102.6 | -3.0 |
| Precipitation $Y$ | 0.0002 | -0.0002 | 0.0005 | 5000 | 55.6 | 82.7 | 0.7 |
| **(Intercept of $X$)** | 3.4115 | 2.7572 | 4.4384 | 5000 | 55.6 | 58.4 | 5.7 |
| **Dist. to road $X$** | -0.0002 | -0.0004 | -0.0001 | 5000 | 55.6 | 387.9 | -3.3 |
| Population $X$ | 0.0000 | -0.0001 | 0.0002 | 5000 | 55.6 | 437.5 | -0.3 |
| $\sigma_Y^2$ | 31.8726 | 21.3638 | 44.6661 | 5000 | - | 8.2 | -3.5 |
| $\sigma_X^2$ | 6.8778 | 4.3181 | 15.4775 | 5000 | - | 5.1 | 2.2 |
| $\text{corr}_{Y,X}$ | 0.972 | 0.906 | 0.994 | - | - | - | - |

### Appendix B.2. Maps of posterior variables for the presence of Pines

**Figure B.6:**
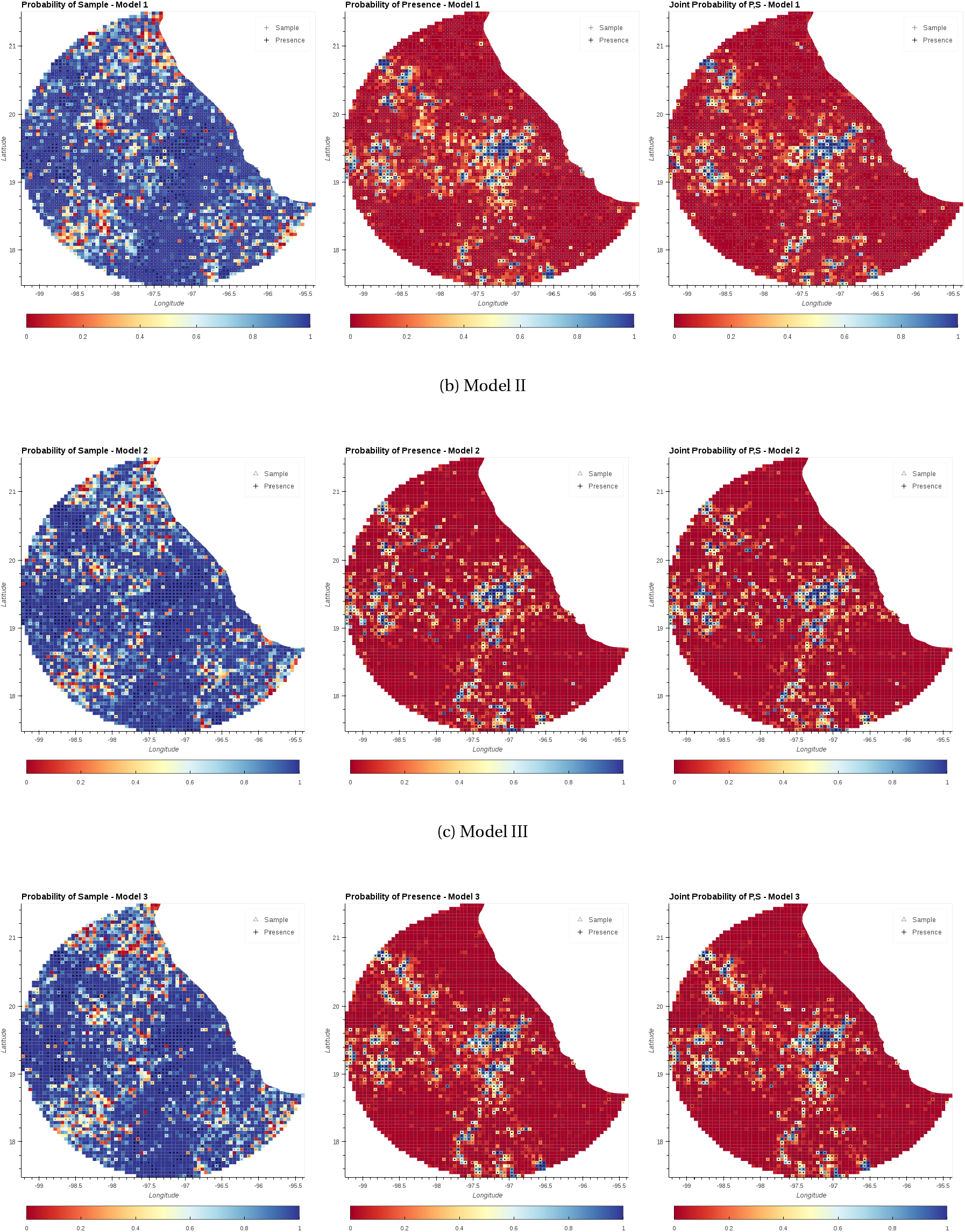
Mean probability and 95% C.I. for Presence, Sample, and Joint presence and sample for Models I, II and III predicting presence of Pines (Class: Pinopsida) using Plants (Kingdom: Plantae) as sample.

**Figure B.7:**
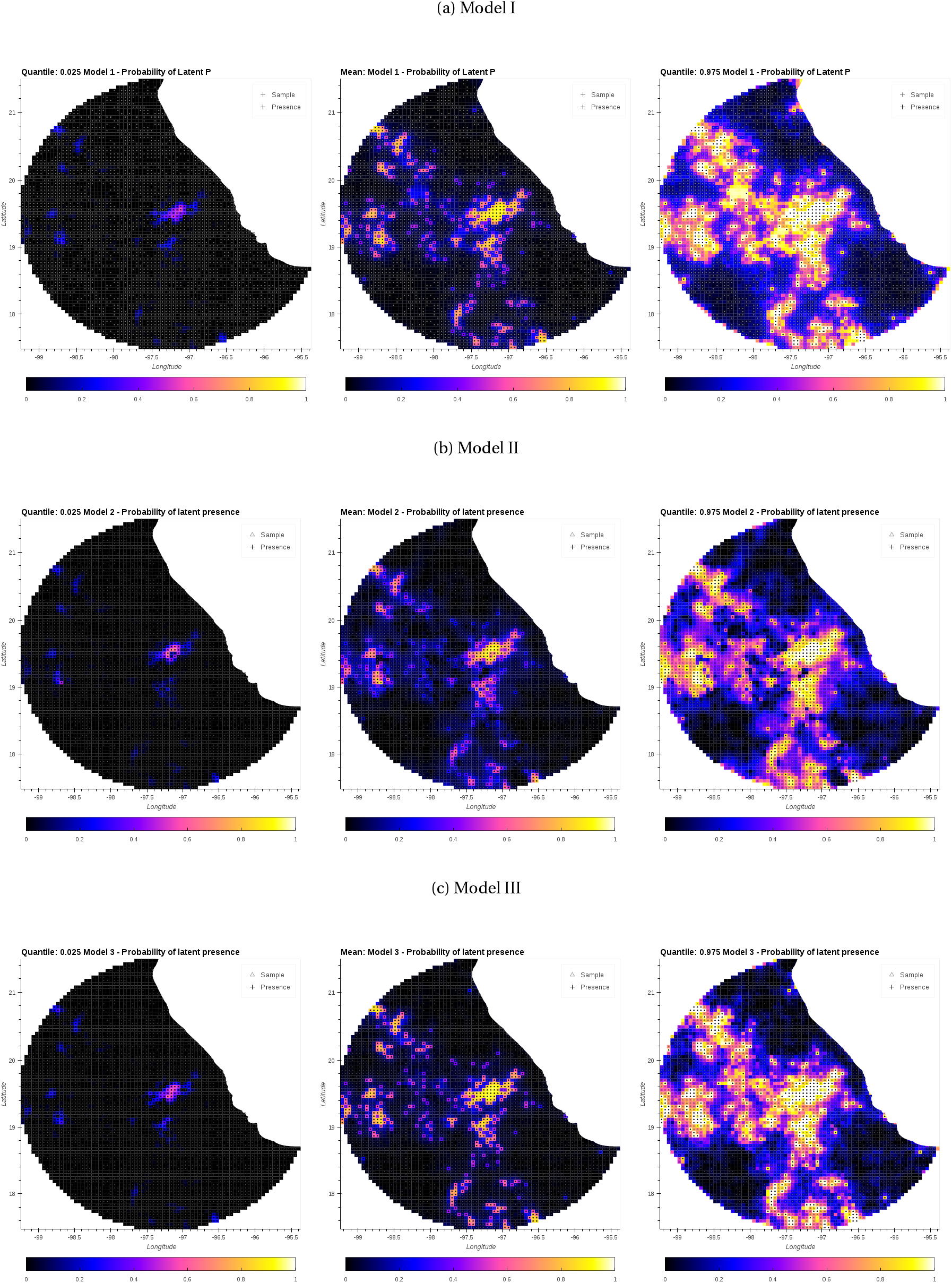
Latent variable *P_Y_* (Presence) for Models I, II and III predicting presence of Pines. The central column corresponds to the mean value. The columns on the left and right correspond to quantiles: 0.025 and 0.975, respectively.

**Figure B.8:**
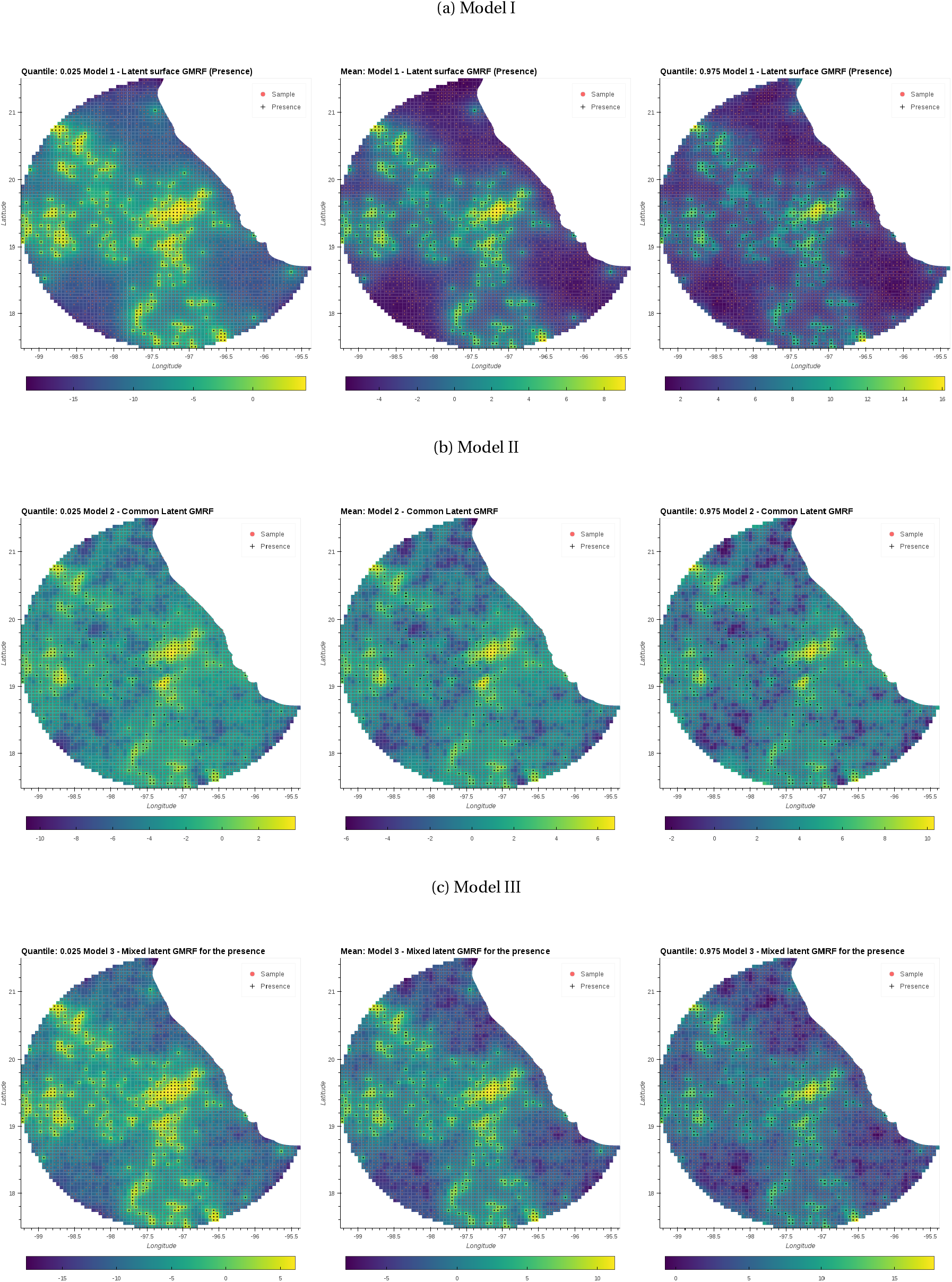
Spatial random effect *S_Y_*. The Gaussian Markov random field (GMRF) corresponding to the latent variable *P_Y_* (Presence) for Models I, II and III predicting presence of Pines. The central column corresponds to the mean value, The column on the left and right corresponds to quantiles: 0.025 and 0.975, respectively.

**Figure B.9:**
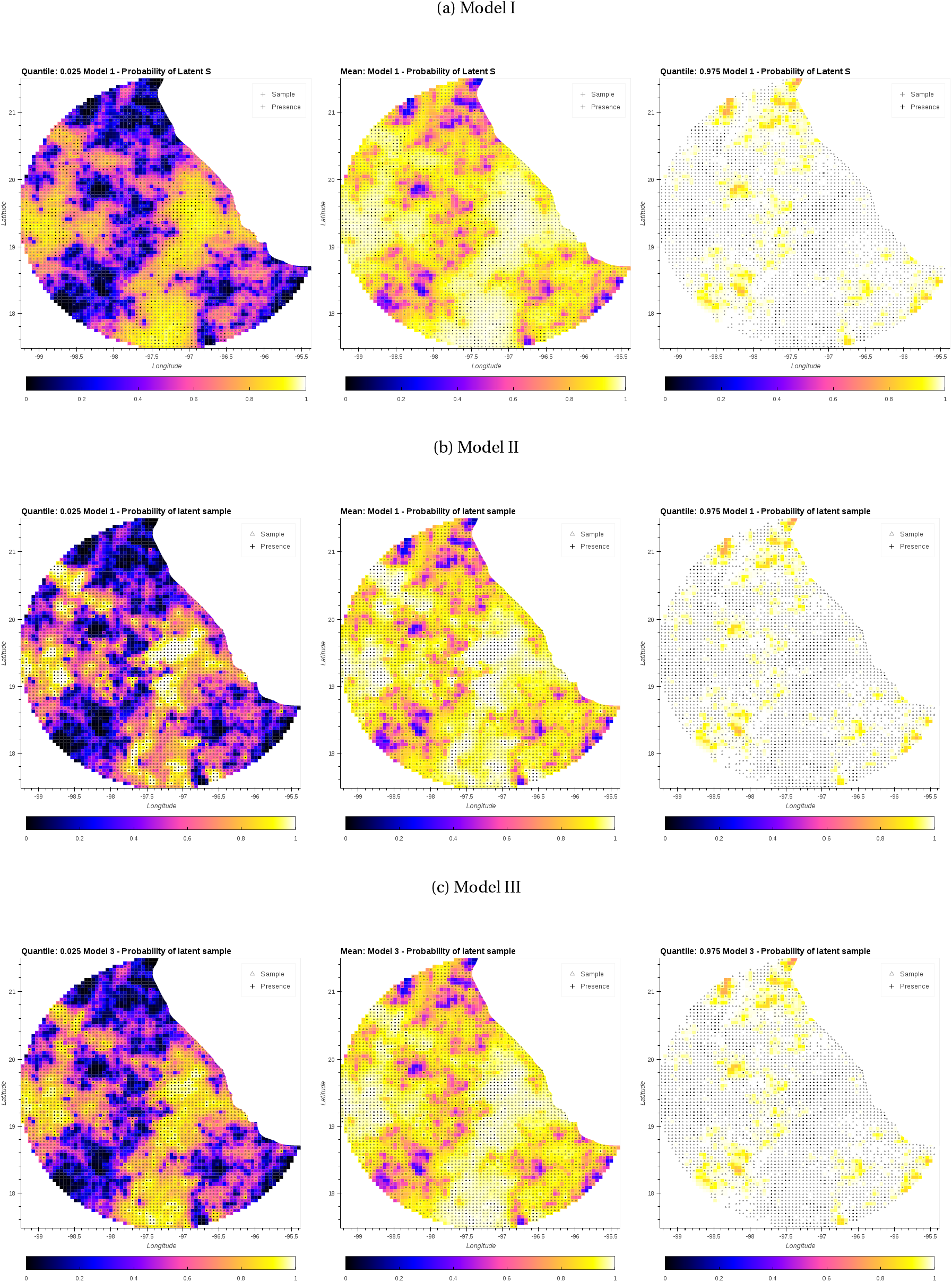
Latent variable *P_X_* (Sample) for Models I, II and III predicting presence of Pines using all plants as sample. The central column corresponds to the mean value. The columns on the left and right correspond to quantiles: 0.025 and 0.975, respectively.

**Figure B.10:**
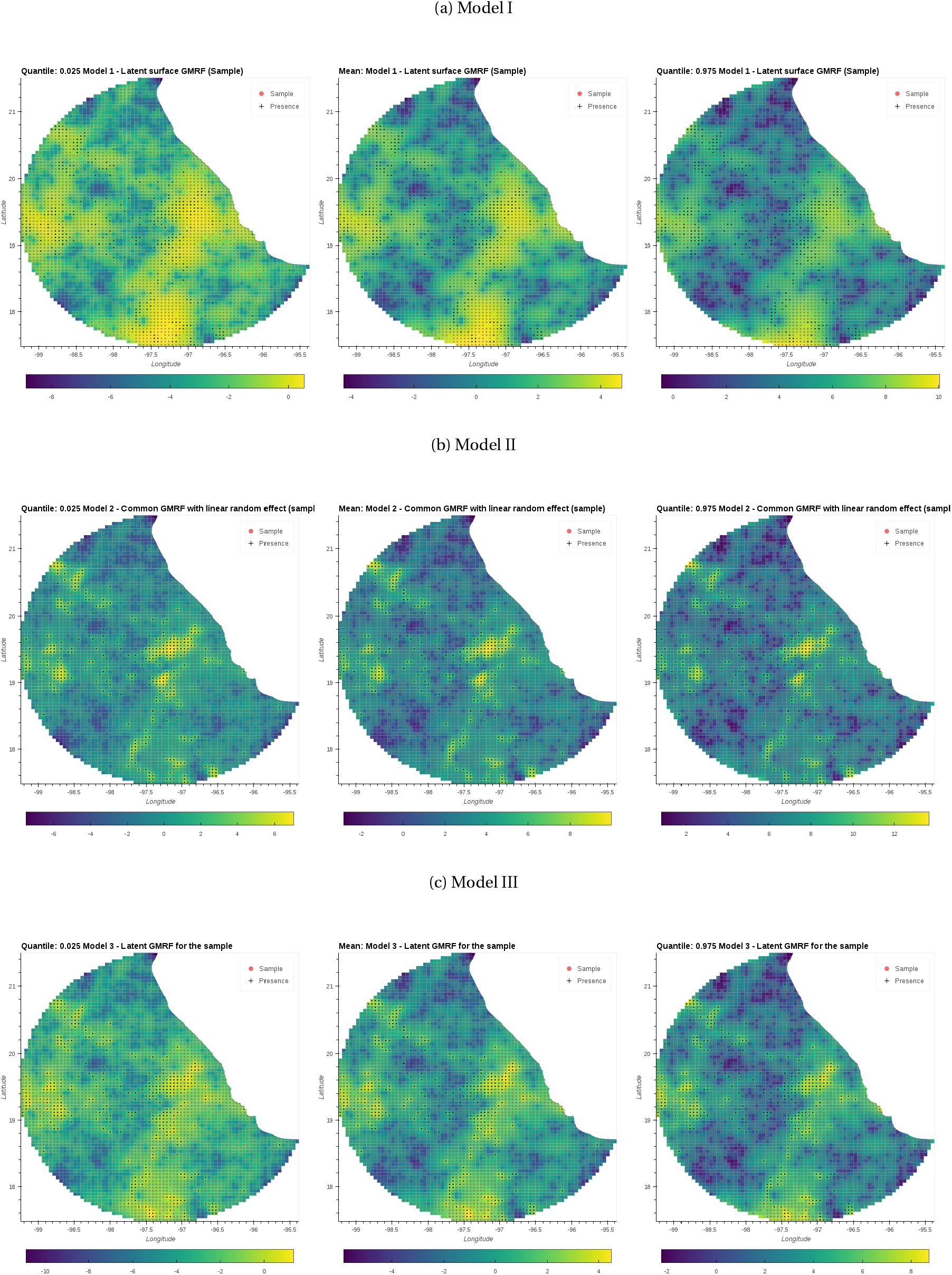
Spatial random effect *S_X_*. The Gaussian Markov random field (GMRF) corresponding to the latent variable *S_X_* (Sample) for Models I, II and III predicting presence of Pines. The central column corresponds to the mean value. The column on the left and right corresponds to quantiles: 0.025 and 0.975, respectively.

**Figure B.11:**
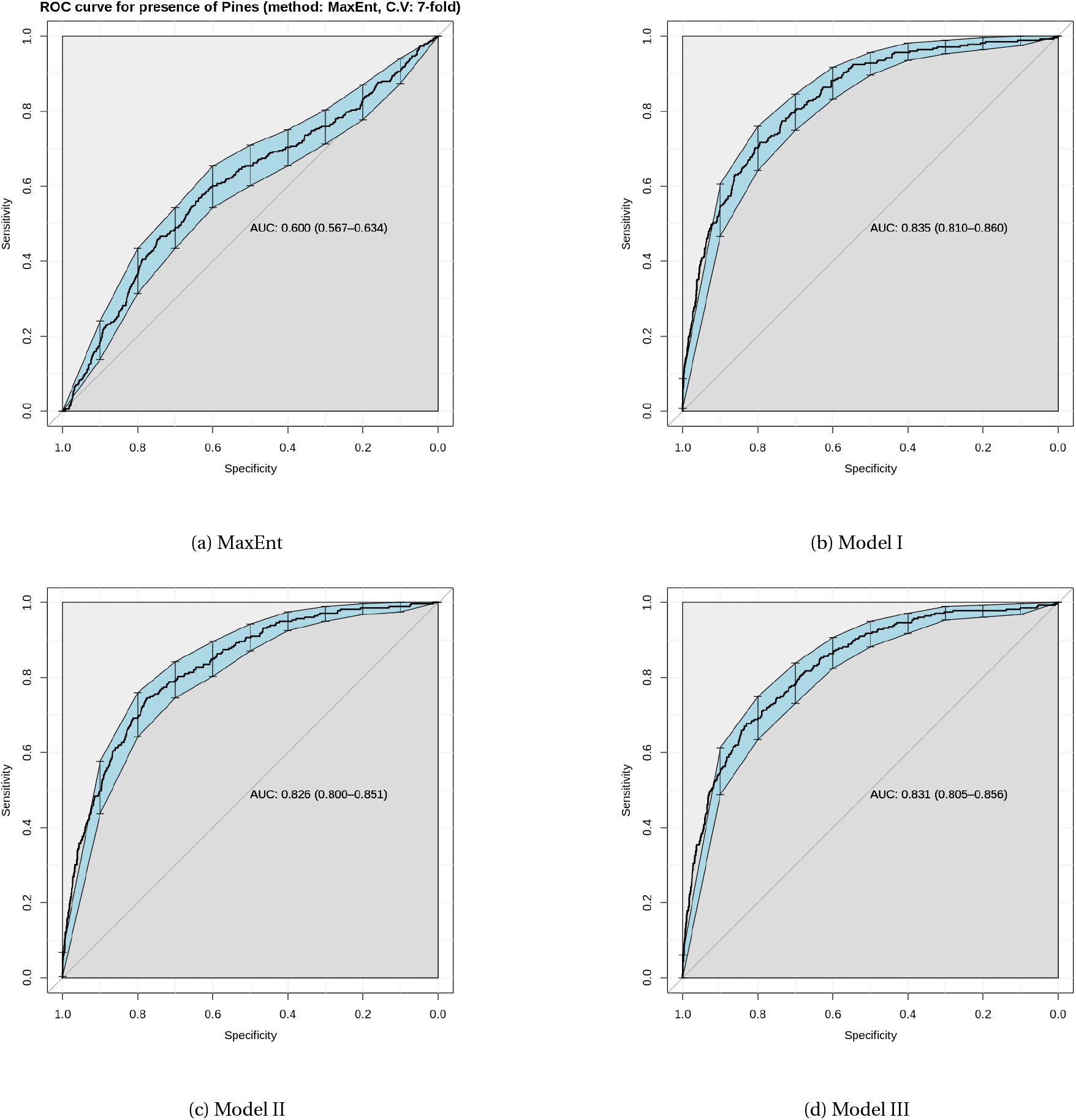
Area under the receiver operating characteristic curve (AUC-ROC) for the different models of Pines. The three models (b,c and d) perform significantly better than MaxEnt.

## Appendix C. Estimates for the predicted presence of tyranids using birds records as sample

**Table C.4:**
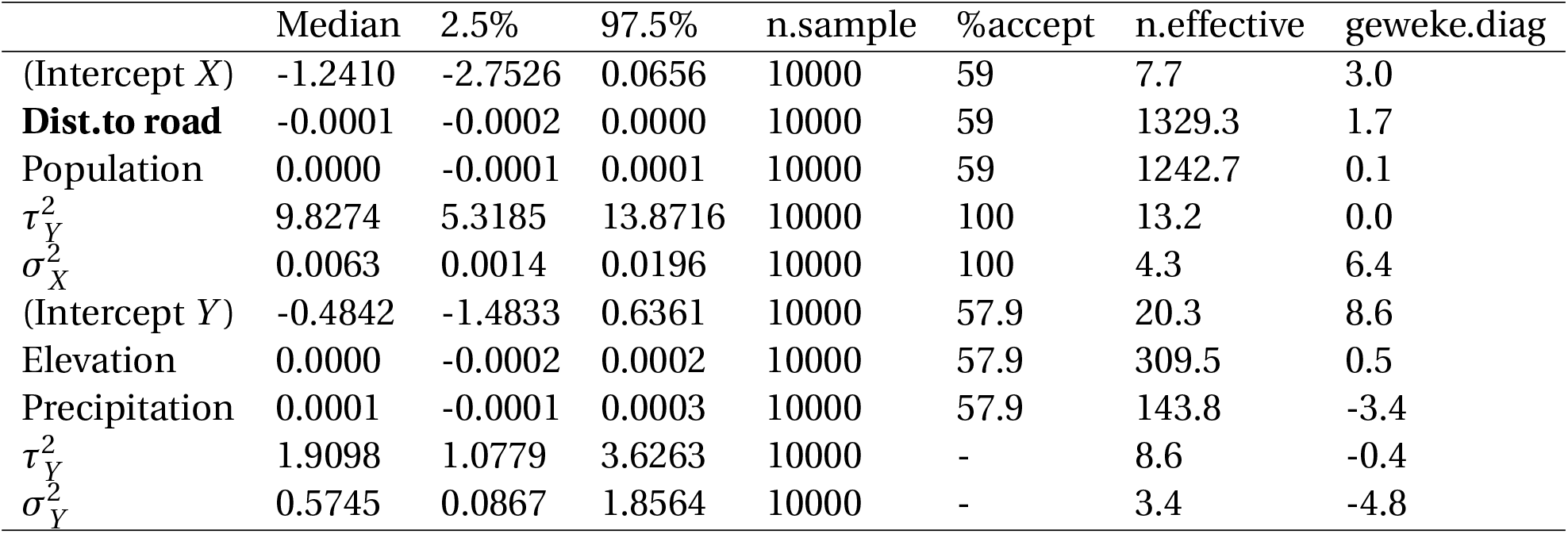
Posterior summaries of all the parameters in model I with the associated 95% credible intervals for the example of flycatchers. Parameters 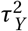 and 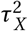 correspond to the variance of the spatial effects of the presence and the sample process (*S_Y_* and *S_X_*) respectively. Likewise, 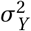 and 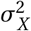 correspond to the variance of the unstructured processes *Z_Y_* and *Z_X_* respectively. Significant parameters for the fixed effect are shown in **bold**. For further information see section: 3

|  | Median | 2.5% | 97.5% | n.sample | %accept | n.effective | geweke.diag |
| --- | --- | --- | --- | --- | --- | --- | --- |
| (Intercept X) | -1.2410 | -2.7526 | 0.0656 | 10000 | 59 | 7.7 | 3.0 |
| **Dist.to road** | -0.0001 | -0.0002 | 0.0000 | 10000 | 59 | 1329.3 | 1.7 |
| Population | 0.0000 | -0.0001 | 0.0001 | 10000 | 59 | 1242.7 | 0.1 |
| $\tau_Y^2$ | 9.8274 | 5.3185 | 13.8716 | 10000 | 100 | 13.2 | 0.0 |
| $\sigma_X^2$ | 0.0063 | 0.0014 | 0.0196 | 10000 | 100 | 4.3 | 6.4 |
| (Intercept Y) | -0.4842 | -1.4833 | 0.6361 | 10000 | 57.9 | 20.3 | 8.6 |
| Elevation | 0.0000 | -0.0002 | 0.0002 | 10000 | 57.9 | 309.5 | 0.5 |
| Precipitation | 0.0001 | -0.0001 | 0.0003 | 10000 | 57.9 | 143.8 | -3.4 |
| $\tau_Y^2$ | 1.9098 | 1.0779 | 3.6263 | 10000 | - | 8.6 | -0.4 |
| $\sigma_Y^2$ | 0.5745 | 0.0867 | 1.8564 | 10000 | - | 3.4 | -4.8 |

**Table C.5:** Posterior summaries of all the parameters in Model II with the associated 95% credible intervals for the example of flycatchers. The parameter *τ*^2^ represents the variance of the common spatial effect. Parameters *σ*^2^ and *σ*^2^ correspond to the variance of the unstructured process *Z_Y_* and *Z_X_*. Significant parameters for the fixed effect are shown in **bold**. For further information see section: 3

|  | Median | 2.5% | 97.5% | n.sample | %accept | n.effective | Geweke.diag |
| --- | --- | --- | --- | --- | --- | --- | --- |
| (Intercept) | -1.6937 | -2.1358 | -1.3629 | 10000 | 47.6 | 68.7 | 4.7 |
| Dist to road | -0.0001 | -0.0002 | 0.0001 | 10000 | 47.6 | 443.7 | -0.8 |
| Population | 0.0000 | -0.0001 | 0.0001 | 10000 | 47.6 | 300.6 | -1.4 |
| Elevation | -0.0001 | -0.0003 | 0.0001 | 10000 | 47.6 | 175.3 | 1.6 |
| Precipitation | 0.0000 | -0.0001 | 0.0002 | 10000 | 47.6 | 192.1 | 2.4 |
| $\tau^2$ | 10.1800 | 7.3033 | 14.9518 | 10000 | - | 18.8 | -3.8 |
| $\sigma^2$ | 0.0089 | 0.0022 | 0.0829 | 10000 | - | 1552.6 | 0.4 |

**Table C.6:** Posterior summaries of all the parameters in Model III with the associated 95% credible intervals for the example of flycatchers. Parameters 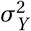 and 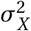 correspond to the variance for the presence (*Y*) and the sample (*X*). The term corr*_x, y_* indicates the correlation between these two processes. Significant parameters for the fixed effect are shown in **bold**. For further information see section: 3

|  | Median | 2.5% | 97.5% | n.sample | %accept | n.effective | Geweke.diag |
| --- | --- | --- | --- | --- | --- | --- | --- |
| (Intercept Y) | -0.9374 | -1.6520 | -0.2057 | 5000 | 53.3 | 110.0 | 1.0 |
| Elevation | 0.0000 | -0.0002 | 0.0002 | 5000 | 53.3 | 88.5 | -1.2 |
| Precipitation | 0.0001 | -0.0001 | 0.0003 | 5000 | 53.3 | 150.2 | -2.0 |
| (Intercept X) | -1.4153 | -1.9346 | -0.9441 | 5000 | 53.3 | 85.2 | 0.4 |
| **Dist. to road** | -0.0001 | -0.0002 | 0.0000 | 5000 | 53.3 | 523.5 | 0.5 |
| Population | 0.0000 | -0.0001 | 0.0001 | 5000 | 53.3 | 232.1 | -1.0 |
| $\sigma_Y^2$ | 3.5179 | 2.7614 | 6.0832 | 5000 | - | 5.6 | -0.7 |
| $\sigma_X^2$ | 7.3840 | 5.9431 | 12.1276 | 5000 | - | 7.1 | -0.6 |
| $\text{corr}_{Y,X}$ | - | - | - | - | - | - | - |

### Appendix C.1. Maps of posterior probabilities for Tyranids

**Figure C.12:**
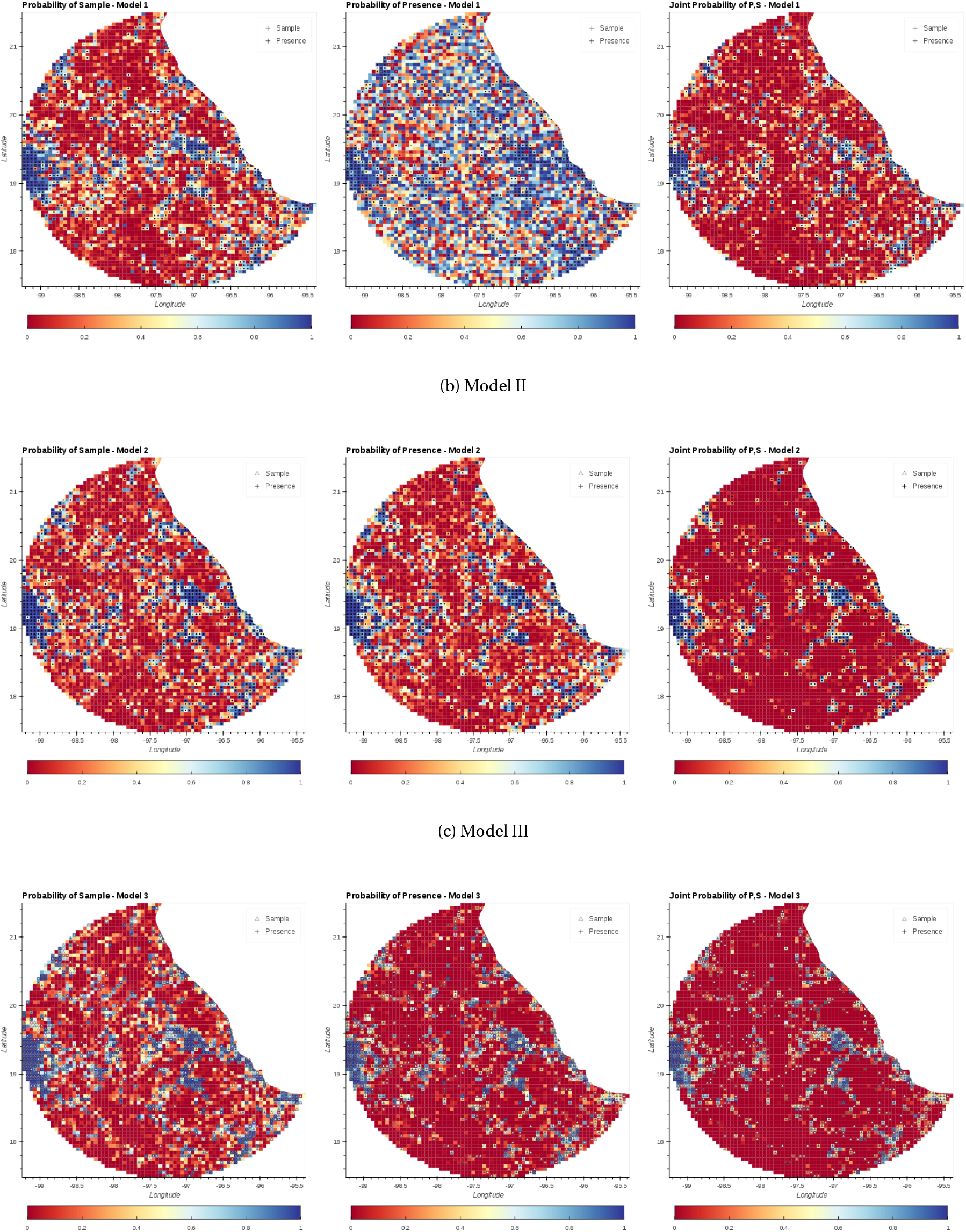
Mean probability and 95% C.I. for Presence, Sample, and Joint presence and sample for Models I, II and III predicting presence of flycatchers (Family: Tyrannidae) using birds (Class: Aves) as sample.

**Figure C.13:**
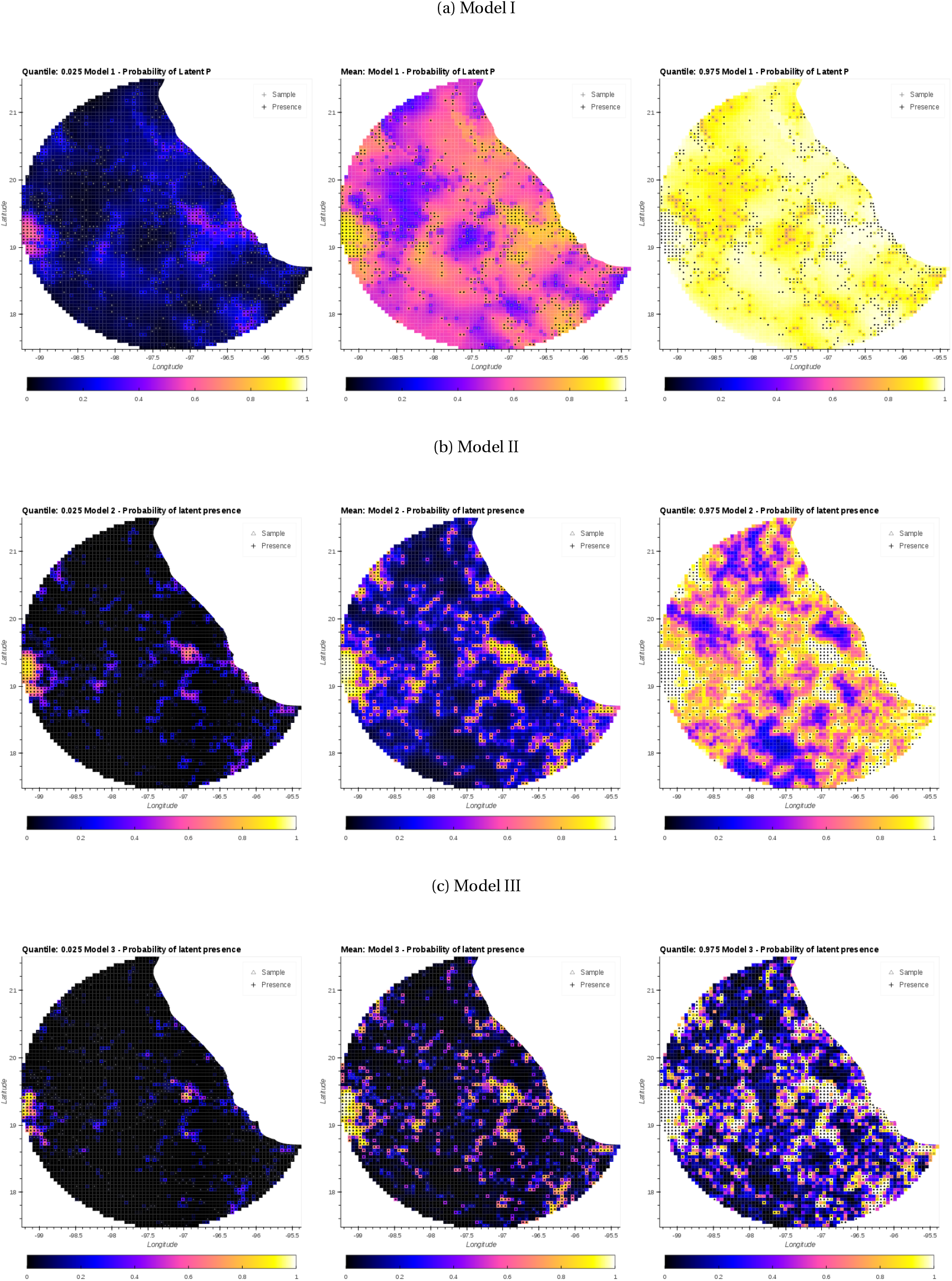
Latent variable *P_Y_* (Presence) for Models I, II and III predicting presence of flycatchers (Family: Tyrannidae). The central column corresponds to the mean value. The columns on the left and right correspond to quantiles: 0.025 and 0.975, respectively.

**Figure C.14:**
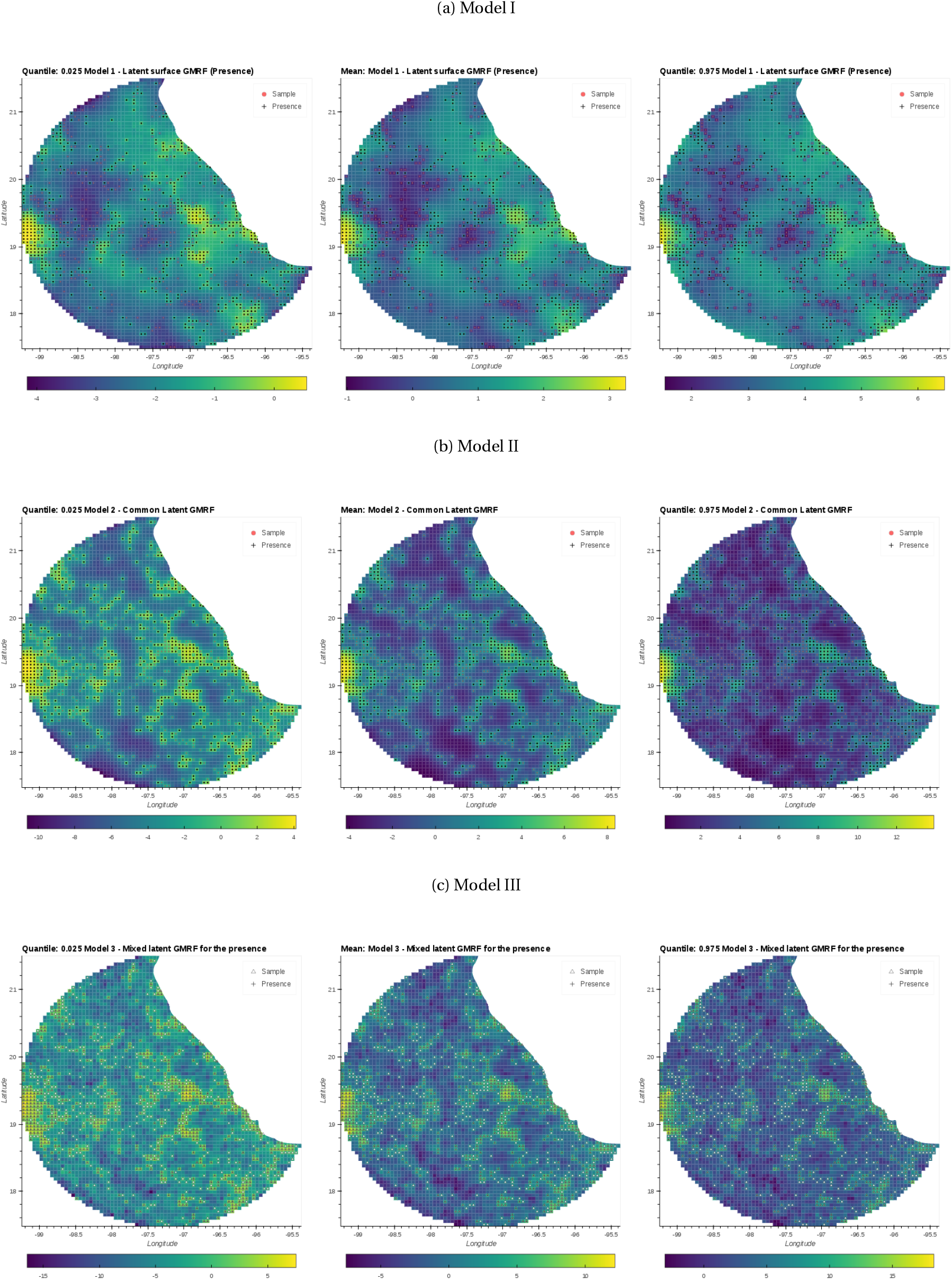
Spatial random effect *S_Y_*. The Gaussian Markov random field (GMRF) corresponding to the latent variable *P_Y_* (Presence) for Models I, II and III predicting presence of flycatchers (Family: Tyrannidae). The central column corresponds to the mean value. The columns on the left and right correspond to quantiles: 0.025 and 0.975, respectively.

**Figure C.15:**
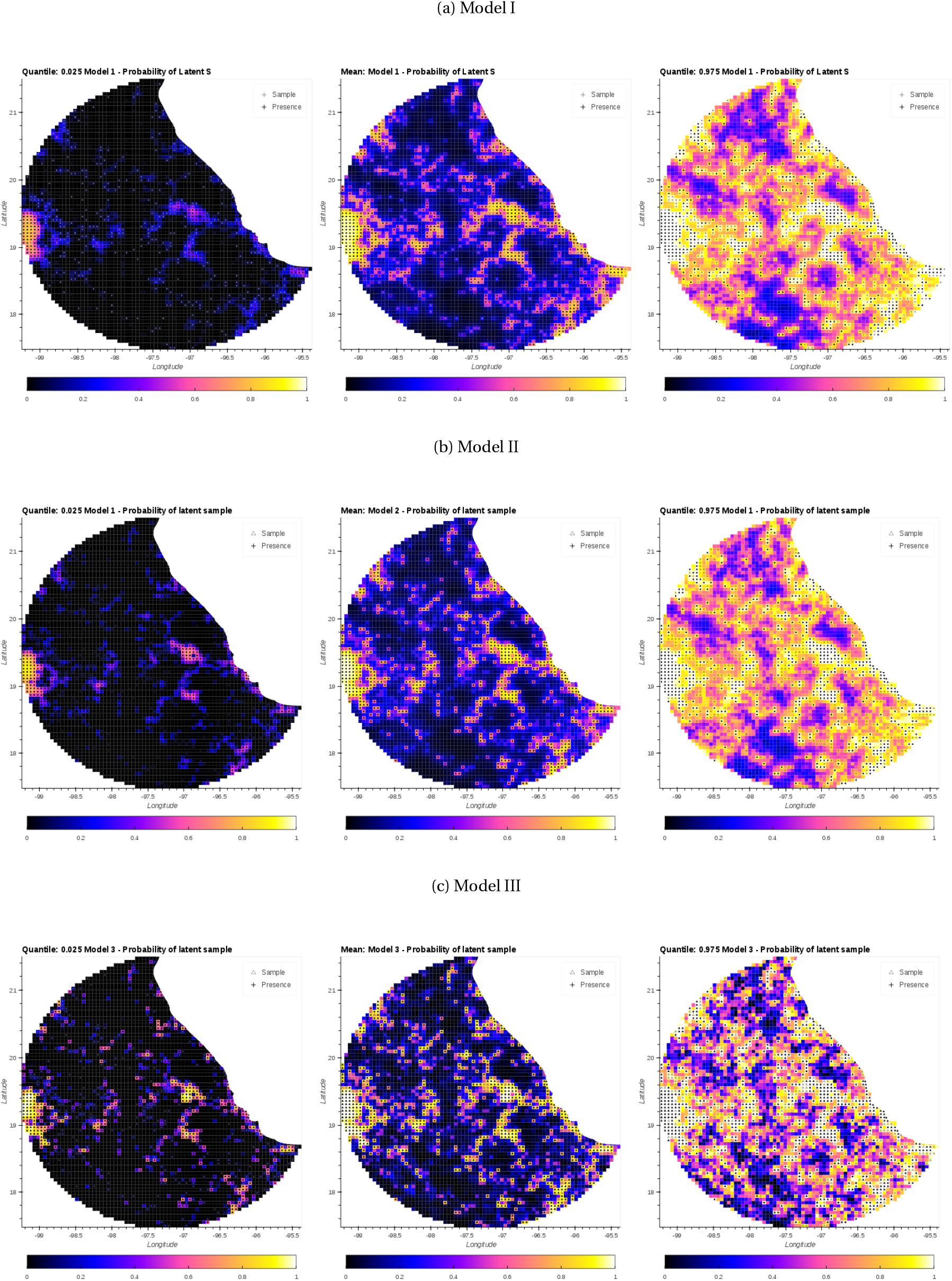
Latent variable *P_X_* (Sample) for Models I, II and III predicting presence of flycatchers (Tyrannidae) using all birds as sample. The central column corresponds to the mean value. The columns on the left and right correspond to quantiles: 0.025 and 0.975, respectively.

**Figure C.16:**
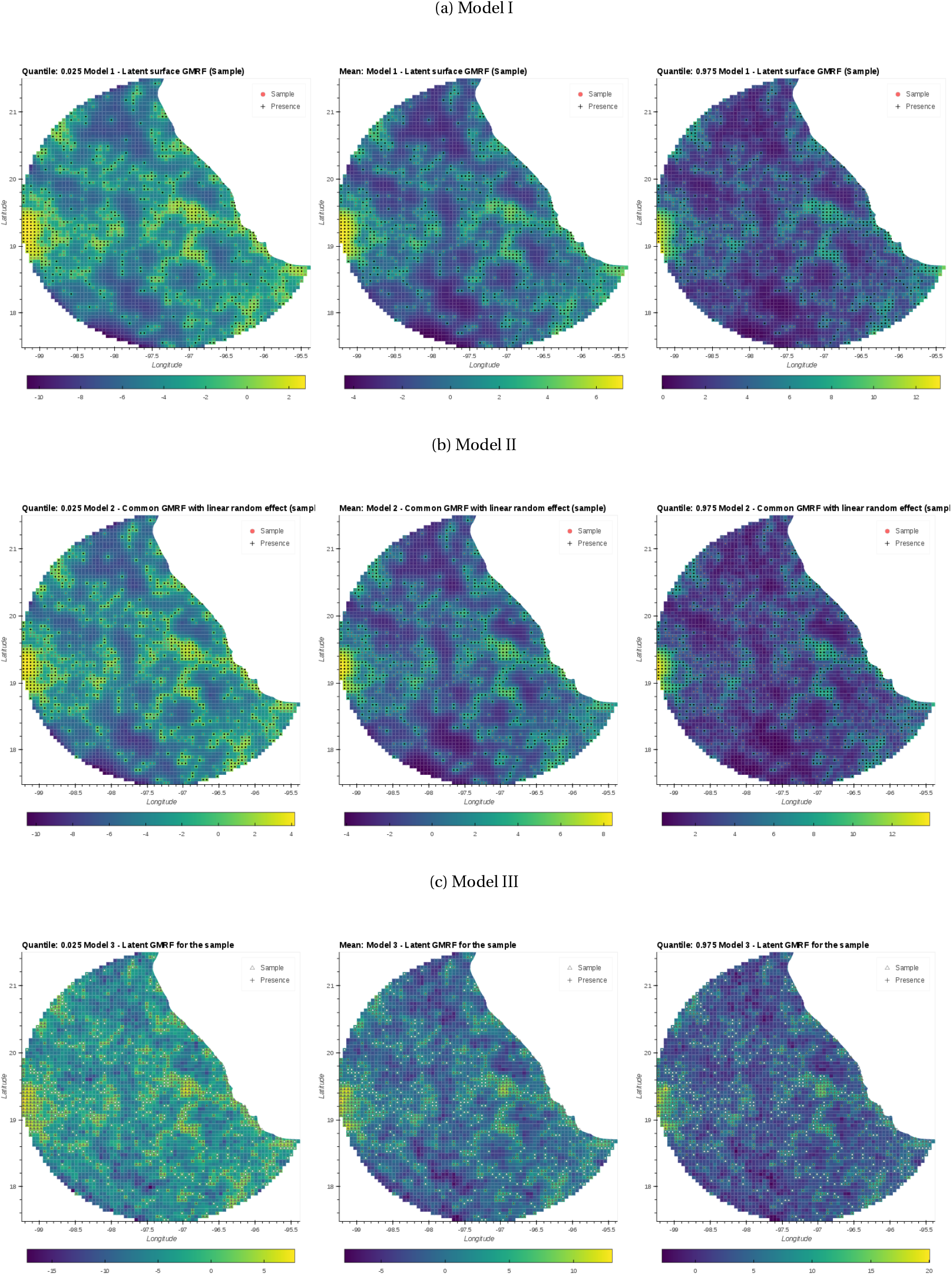
Spatial random effect *S_X_*. The Gaussian Markov random field (GMRF) corresponding to the latent variable *P_X_* (Sample) for Models I, II and III predicting presence of flycatchers (Tyrannidae). The central column corresponds to the mean value. The columns on the left and right correspond to quantiles: 0.025 and 0.975, respectively.

**Figure C.17:**
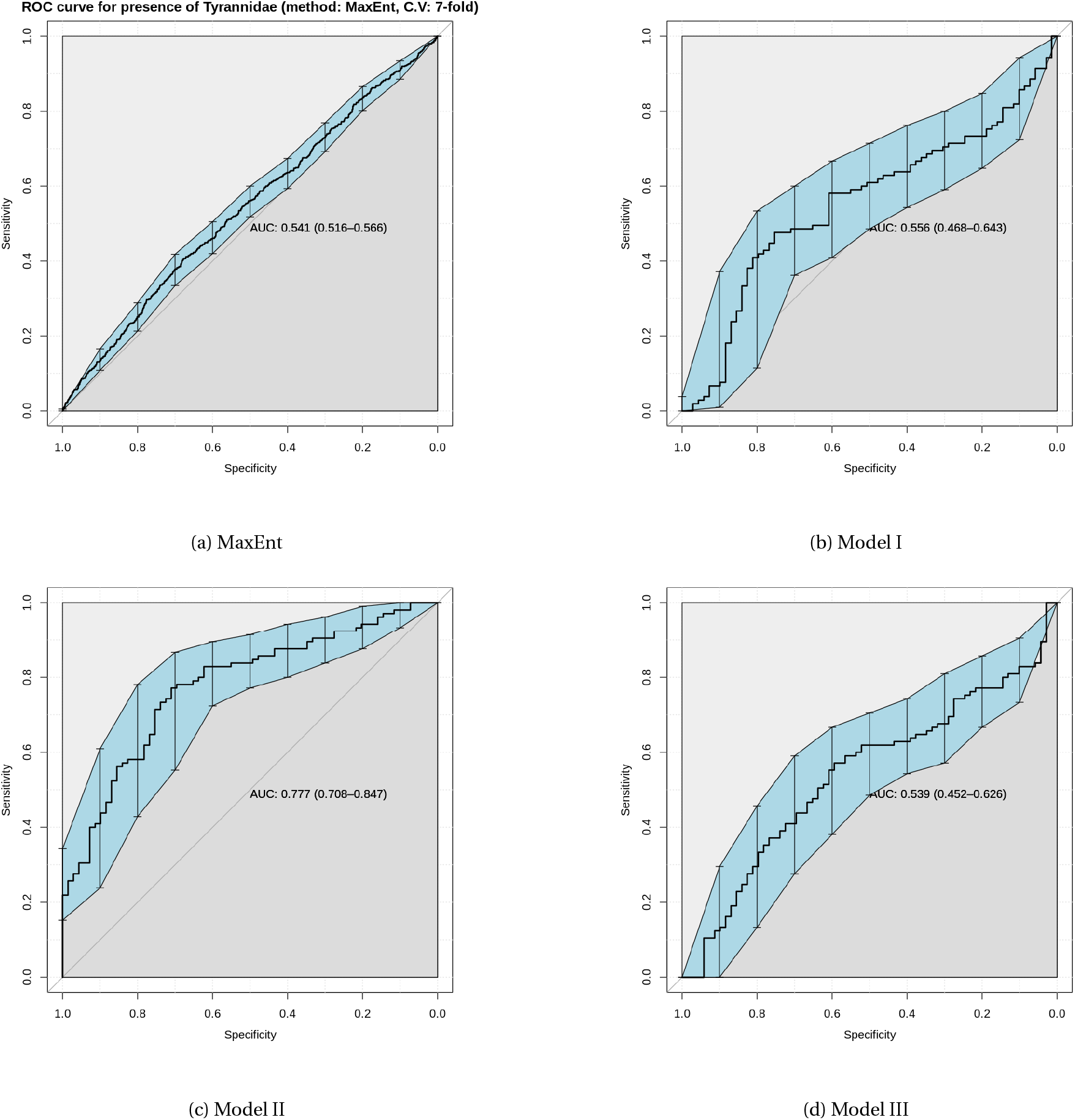
Area under the receiver operating characteristic curve (AUC-ROC) for MaxEnt and models I, II and III of flycatchers. MaxEnt and models I and III achieved low AUC. Although, on average models I and III out performed MaxEnt, their variances show that these models are not appropriate when the proportion of missing data is significantly higher than the presences. See the discussion section for a more detail explanation.

